# Modulating Sleep Oscillations Through Targeted Memory Reactivation: Linking Functional Dynamics to Brain Microstructure

**DOI:** 10.64898/2026.10.13.758910

**Authors:** Whitney Stee, Michael-Christopher Foti, Antoine Legouhy, Michele Guerreri, Hui Zhang, Jean-Marc Lina, Philippe Peigneux

## Abstract

**Background:** Sleep-dependent memory consolidation reputedly rely on the coordinated activity of sleep oscillations, which may facilitate the reactivation and redistribution of newly acquired memories. Targeted memory reactivation (TMR) during sleep can experimentally modulate these processes, although effects remain variable. Moreover, how learning-related microstructural plasticity relates to sleep oscillatory activity and spindle-related network dynamics remains poorly understood. We investigated whether auditory TMR modulates sleep spindles, slow oscillations (SOs) and their co-occurrence, whether these changes relate to motor memory consolidation and diffusion MRI (dMRI) measures of brain microstructure, and whether learning and TMR alter the evolution of cortico-cortical connectivity during spindle development.

**Methods:** Sixty healthy young adults underwent five dMRI sessions before and after motor sequence learning (Day 1), after a post-training night of regular sleep (RS) or TMR (Day 2), and before and after relearning (Day 5). Baseline and experimental nights were recorded with 256-channel high-density EEG. Oscillation densities were analyzed using mixed-design ANOVAs and related to behavioral and dMRI changes using Pearson or Spearman correlations. Intra-spindle connectivity between 148 source-reconstructed cortical regions was compared between spindle onset and end, across nights and between groups using Network-Based Statistics.

**Results:** Spindle density, SO density and SO–spindle co-occurrence remained stable across nights in the RS group but decreased after TMR. Higher spindle density and co-occurrence were associated with greater overnight performance gains, and all oscillatory measures were associated with learning-related dMRI changes in subcortical regions, notably striatum and thalamus. Connectivity increased from the beginning to the end of the spindle duration during the baseline night in both groups, but during the experimental night only in the RS group.

**Conclusions:** Motor learning is associated with coordinated changes in sleep oscillatory activity, within-spindle cortical connectivity and brain microstructure, supporting a dynamic interaction between learning-induced plasticity and sleep-dependent network reorganization. Under the present stimulation conditions, TMR did not enhance these processes but instead reduced sleep oscillatory activity and attenuated spindle-related connectivity dynamics, underscoring the importance of precisely targeting endogenous sleep rhythms when using TMR.

## **1.** Background

### 1.1. Sleep oscillations and memory consolidation

Spontaneous reactivation of learning-related neural patterns during post-learning sleep has been associated with subsequent memory consolidation and retrieval [1]. In animals, selective disruption of the reactivation process using optogenetic approaches impairs memory recall, suggesting causal evidence for its role in consolidation [2]. In humans, spontaneous replay of motor memories has similarly been observed during sleep, with reactivation within the striato-cerebello-cortical network time-locked to spindle occurrence [3, 4]. Moreover, the magnitude of this reactivation was associated with subsequent improvements in motor performance [5]. These findings suggest that sleep-dependent motor memory consolidation involves a coordinated reactivation of distributed neural systems, with sleep spindles potentially providing a temporal window for the interaction and functional integration of regions including the hippocampus, striatum, thalamus, and motor-related cortical areas [3, 4, 6, 7].

In addition to sleep spindles, cortical slow oscillations (SOs) also appear to play a crucial role in memory consolidation [8, 9]. SOs are thought to provide a temporal framework for the coordinated reactivation of learning-related neural populations and the local reprocessing of distinct memory components [7, 10]. Through this coordination, SOs may facilitate the interaction between distributed brain regions and spindle activity, thereby promoting the integration and consolidation of newly acquired memory representations [7].

Increasing evidence suggests that memory consolidation depends not only on the occurrence of SOs or spindles individually, but also on their precise temporal coordination [11–13]. Spindles, particularly fast spindles (12–15 Hz), preferentially coincide with the SO upstate, corresponding to cortical depolarization, whereas slower spindles (9–12 Hz) are more likely to occur during the transition from the up- to down-state, [14–16]. Higher SO–spindle coupling has been associated with enhanced sleep-dependent memory consolidation both in the declarative [17–21] and motor memory domains [20, 22]. Conversely, reduced temporal coordination between SOs and spindles, as observed in children [17, 22] and older adults [19, 23] has been associated with diminished overnight memory gains. Consistently, SO-coupled spindles have been shown to predict memory consolidation more strongly than uncoupled spindles [11, 24]. Experimental manipulations using electrical [25, 26] and acoustic stimulation [27, 28], as well as GABAergic drug administration [10], further demonstrated that SO–spindle coupling can be enhanced, although effects on memory have been inconsistent [10, 25, 26, 28]. Importantly, recent evidence linked the precision of SO–spindle coupling to the strength of memory reactivation during sleep, with stronger post-learning reactivation predicting greater subsequent memory consolidation [29]. Together, these findings suggest that the temporal coordination of SOs and spindles may provide a key mechanism through which newly encoded information is reactivated and integrated during sleep.

### 1.2. Targeted memory reactivation

Expanding on the role of sleep-related reactivation in memory consolidation, Targeted Memory Reactivation (TMR) was introduced as a method to selectively reactivate recently encoded memory traces during sleep [30] (for reviews, see [31, 32]). TMR involves presenting sensory cues during sleep that were previously associated with specific information during wakeful learning. Across studies, TMR was shown to selectively enhance memory for cued as compared with non-cued material [33], with benefits across multiple domains including verbal [34–38], spatial [39, 40], fear extinction [41], and motor memory [42–45]. However, meta-analytic evidence indicates that these effects are generally small to moderate and may vary substantially across tasks and memory domains [33].

The impact of TMR appears to be closely linked to its ability to modulate sleep oscillatory activity. Auditory cueing during sleep was shown to increase spindle activity [42, 46], particularly over parietal regions [42], with greater spindle responses associated with stronger behavioral benefits [42, 43]. TMR was also reported to increase the occurrence of SOs time-locked to spindles, potentially enhancing their temporal coordination [44, 46]. However, spindle modulation is not consistently observed across studies even when using similar stimulation protocols [44], suggesting that the effects of TMR critically depend on the timing and parameters of cue presentation. Thus, although TMR can enhance memory consolidation, its efficacy may depend on its ability to engage, rather than disrupt, the endogenous oscillatory dynamics supporting memory reactivation.

### 1.3. Microstructure & sleep oscillations

Sleep oscillation features such as sleep spindles [47–49], slow-wave activity (SWA) [50], and the slope of SOs [51] have been consistently associated with brain plasticity processes and its structural properties. These include inter-individual differences in grey-matter (GM) volume [47, 49, 50, 52], cortical GM thickness [53], and white matter (WM) diffusion properties [50]. For example, steeper SO rising slopes and greater spindle power have been associated with higher axial diffusivity in WM tracts across frontal, temporal, and subcortical regions [47]. Similarly, WM microstructure was linked to spindle density and learning-related memory gains, suggesting that structural properties may contribute to spindle-dependent memory consolidation [48]. Regional GM characteristics were also associated with sleep oscillatory activity, the hippocampal GM volume being related to spindle expression, and hypothalamic and medial prefrontal GM volume related to SO activity [52]. Together, these findings suggest that sleep oscillations are not only functional signatures of neural network activity but are also shaped by the structural properties of their underlying neural substrate. Conversely, sleep spindles are thought to promote cellular processes underlying neural plasticity [54], raising the possibility of a bidirectional relationship between sleep oscillations and brain structure. Whether experimentally manipulating sleep oscillations through TMR can influence learning-related microstructural changes, however, remains unknown.

### 1.4. Intra-spindle connectivity dynamics

Beyond their occurrence and coupling with SOs, the temporal dynamics of neural communication within individual spindles may provide additional insights into their role in memory processing. Dynamic changes in brain connectivity were observed across the 1–2 s duration of sleep spindles, characterized by a transition from predominantly localized, intra-hemispheric connectivity during the early spindle phase toward a more widespread inter- hemispheric communication during the later phase [55]. Such dynamics may underlie the progressive coordination of distributed neural populations during spindle events. Whether this intra-spindle connectivity evolution is influenced by recent learning, and whether it can subsequently be modulated by TMR, remains largely unexplored.

To sum up, previous work indicates that sleep-dependent memory consolidation involves the coordinated activity of spindles and SOs, their precise temporal coupling, and the progressive interaction of distributed neural networks. TMR provides a means of experimentally manipulating these processes, but its effects on endogenous oscillatory dynamics and spindle-related network communication remain incompletely understood. Moreover, the relationship between these sleep oscillations and learning-induced changes in brain microstructure remains unclear.

In this framework, we investigated in the present study how TMR influences sleep oscillations and their relationship with motor memory consolidation and learning-related microstructural changes. Specifically, we examined (1) whether TMR modulated spindle and SO density, SO– spindle co-occurrence, and (2) whether these oscillatory features were associated with overnight changes in motor performance and learning-related dMRI measures. We hypothesized that higher spindle and SO–spindle activity would be associated with more efficient memory consolidation and greater learning-related microstructural changes. We further expected TMR to enhance spindle-related activity and potentially increase SO–spindle coordination.

In a second step, we examined the temporal evolution of cortico-cortical connectivity within individual sleep spindles. We first assessed whether recent motor learning modified the progressive change in connectivity from the beginning to the end of spindles, and then tested whether TMR further modulated these post-learning connectivity dynamics. We expected learning to enhance intra-spindle connectivity and investigated whether TMR modulated this effect.

## 2. Methods

### 2.1. Participants

Sixty healthy young adults (30 females) aged 18-29 years (mean age ± SD: 21.58 ± 2.17) participated in this study approved by the Ethics Committee of the Liège University Hospital (approval no. 2020/138). All participants provided written informed consent prior to participation. Eligibility criteria included the absence of neurological or psychiatric conditions, a body mass index (BMI) < 28, and no contraindications to EEG or MRI. Participants exhibited moderate to neutral chronotypes (mean Morningness–Eveningness Questionnaire [56] score ± SD: 54.28 ± 7.10; range: 32–68) and reported good sleep quality (mean Pittsburgh Sleep Quality Index [57] score ± SD: 3.47 ± 1.40; range: 0–6). To minimize confounding factors, musicians and computer scientists (due to potential superior hand motor dexterity from practice), smokers, and individuals who experienced jet lag in the previous three months were excluded. Both right- and left-handed individuals were included (mean Edinburgh

Handedness Inventory [58] score ± SD: 4.85 ± 6.60; range: -10 to 10) since the motor task required bimanual involvement, and structural asymmetries related to handedness are not consistently observed (see [59–61]). Participants were pseudo-randomly assigned to either the Regular Sleep (RS) or the TMR sleep (TMR) group (see below), with each group comprising 15 women and 15 men. This sample is identical to the one used in [62], where additional participants’ characteristics are provided.

### 2.2. Motor Sequence Learning Task

In this study, participants performed a 6-choice version of the Serial Reaction Time Task (SRTT; see Figure 1) coupled with auditory tones, implemented using PsychoPy3 v2020.2.10 (Nottingham, UK). Participants were seated in front of a computer screen and instructed to place the index, middle, and ring fingers of both hands on six response keys corresponding to six horizontally aligned squares displayed on the screen. Their task was to press as quickly and accurately as possible the key corresponding to the square in which a visual cue appeared. Each key press additionally triggered a unique auditory tone associated with its corresponding key (1 = C, 2 = C♯, 3 = D, 4 = D♯, 5 = E, 6 = F), and the next trial began after a 500-ms inter-trial interval. Trials followed either a fixed 12-element sequence (5-3-1-6-2-4-1-5-2-3-6-4) or a pseudo-random order, with the constraint that the same position could not occur on two consecutive trials. Each block comprised 96 key presses, corresponding to 8 repetitions of the 12-element sequence. The initial learning session comprised 30 blocks, including one pseudo- random block (block 26). The following morning, participants completed a retest session consisting of two sequential blocks (blocks 31–32). After three additional nights, participants performed a relearning session comprising 20 blocks (blocks 33–52), with pseudo-random blocks interspersed at blocks 35 and 48. Pseudo-random blocks were included to dissociate sequence-specific learning from sequence-independent motor learning, both of which can contribute to performance improvements [63]. Participants were not informed that the task followed a repeating sequence.

**Figure 1.**
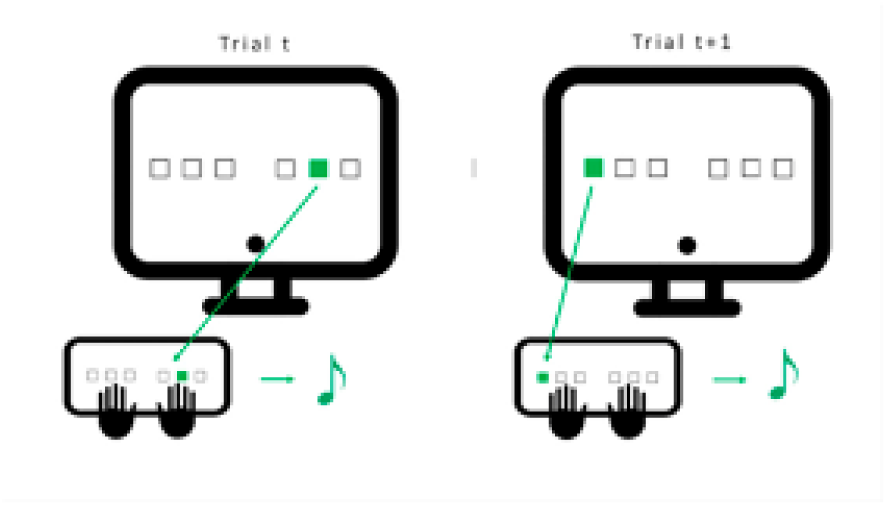
Serial Reaction Time Task (SRTT) Participants sat in front of a computer screen with index, middle, and ring fingers of both hands positioned on six response keys corresponding to six horizontally aligned positions displayed on the screen. Their task was to press the key matching the highlighted position as quickly and accurately as possible. Each key press triggered a unique auditory tone associated with that key, and the next visual cue appeared after a 500-ms delay. Each block comprised 96 trials presented either as a repeating 12-element sequence or in a pseudo-random order.

### 2.3. Experimental Procedure

The procedure is identical to that described in [62], only information relevant to the present analyses is summarized here.

Participants were instructed to abstain from caffeine and other stimulants on testing days and to maintain a regular sleep-wake schedule throughout the study. Sleep schedule regularity was monitored through self-reported sleep logs (using the St. Mary’s Hospital Sleep Questionnaire [64]) and visual inspection of actigraphy data (Actigraph™ wGT3X-BT, Pensacola, FL, USA) recorded from three days before the first testing day until the end of the experiment. To minimize hormonal influences on motor performance, sleep, and memory consolidation, female participants were tested exclusively during the luteal phase of their menstrual cycle [65, 66].

The experimental timeline is illustrated in Figure 2. Three days before the first testing session (Day 1), participants completed a habituation night in the laboratory with 256-channel high-density EEG (hd-EEG). On Day 1 (approximately 16:30), participants underwent a baseline diffusion-weighted MRI scan (DWI1), followed by 30 learning SRTT blocks (for approximately 1 h; see Section 2.2). Thirty minutes after completing the task, a second diffusion-weighted MRI scan (DWI2) was acquired. Participants were then pseudo-randomly assigned to either the RS or TMR group for the post-learning night, while remaining blinded to group allocation. Around 21:30, the hd-EEG cap was fitted, and participants slept in the laboratory for approximately 8–9 h at their habitual bedtime.

**Figure 2.**
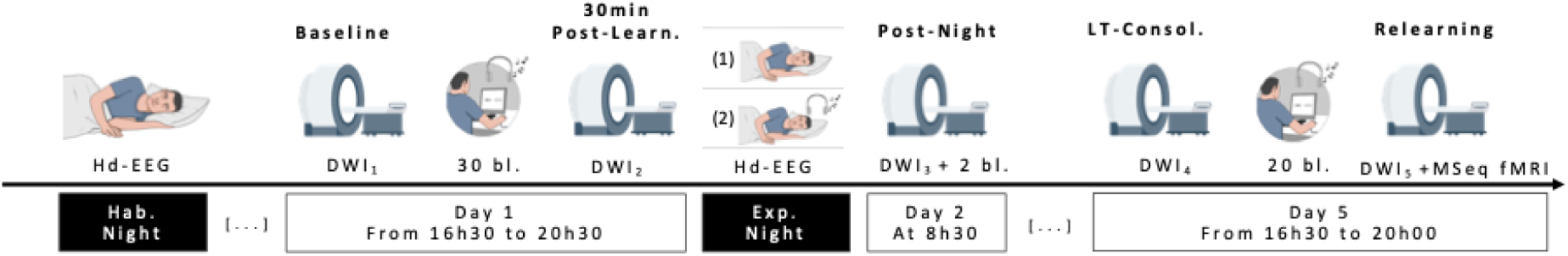
Experimental design Three days before the first testing day, participants completed a habituation night in the laboratory under high -density EEG (hd-EEG) to acclimate to the sleep environment. On Day 1, participants underwent a baseline diffusion MRI scan (DWI1), followed by approximately one hour of motor sequence learning on the Serial Reaction Time Task (SRTT; 30 blocks), during which each response key was paired with a specific auditory tone. Thirty minutes after the learning session, a second diffusion MRI scan (DWI2) was acquired. During the subsequent night, participants either slept without auditory stimulation (RS group) or received learning-related auditory cues during N2 and N3 sleep (TMR group), while hd-EEG was continuously recorded. The following morning (Day 2), participants underwent a third diffusion MRI scan (DWI3) followed by a brief behavioral retest on the SRTT (two blocks). After three nights of regular sleep at home, participants returned to the laboratory on Day 5 for a fourth diffusion MRI scan (DWI4), followed by a 40-minute SRTT relearning session (20 blocks). The experiment concluded with a final diffusion MRI scan (DWI5).

Before sleep, auditory stimulation was individually calibrated to determine the highest sound intensity that participants could comfortably tolerate without being awakened. Participants listened to the task-related tones presented in a random order while the experimenter progressively adjusted the sound level.

During the experimental night, the TMR group received auditory stimulation during stable Non-Rapid Eye Movement sleep stage 3 (N3) in the first sleep cycle and during both Non-Rapid Eye Movement sleep stage 2 (N2) and N3 sleep in the second cycle. Because each response key of the SRTT was paired with a unique auditory tone (Section 2.2), the learned 12-tone sequence ("melody") was replayed during sleep. Each stimulation sequence consisted of twelve 500-ms tones (6 sec in total), followed by a 6-sec silent interval before the next presentation. Auditory stimulation was suspended whenever signs of arousal or transitions to non-target sleep stages were detected and resumed once stable N2 or N3 sleep was re-established.

On Day 2, at least 30 min after awakening (approximately 08:30), participants underwent a third diffusion-weighted MRI scan (DWI3), followed by a brief SRTT retest consisting of two sequential blocks performed outside the scanner. Participants were then sent home and instructed to continue maintaining a regular sleep–wake schedule. On Day 5 (16:30), matching the timing of the initial testing session to control for circadian influences [67, 68], participants underwent a fourth diffusion-weighted MRI scan (DWI4), then completed a 20- block relearning session (for approximately 40 min), and underwent a final diffusion-weighted MRI scan (DWI5) 30 minutes later. To monitor fluctuations in vigilance, participants also completed a 5-min version of the Psychomotor Vigilance Test (PVT) [69] before each SRTT session.

### 2.4. Behavioral data analysis

SRTT performance was evaluated for each block using the Global Performance Index (GPI) [42] ranging from 0 to 1 (with 1 representing ideal performance). The GPI accounts for individual differences in strategy by integrating both accuracy and speed measures. Additional details regarding the behavioral analyses can be found in [62].

### 2.5. High-Density EEG Acquisition and Preprocessing

EEG data were acquired using a 256-channel high-density EEG system (Geodesic EEG System, Electrical Geodesics, Inc.), equipped with HydroCel Geodesic Sensor Nets designed to ensure comprehensive and consistent scalp coverage. Signals were recorded at a sampling rate of 250 Hz with a band-pass filter of 0.1–100 Hz. Electrode impedances were maintained below 70 kΩ at the beginning of each overnight recording session. EEG data was continuously recorded throughout the night, with the timing of acoustic stimulations marked on the recordings.

Sleep scoring was manually performed following American Academy of Sleep Medicine (AASM) guidelines [70] using a subset of electrodes positioned at F3, F4, C3, C4, O1, together with two electrooculography (EOG) and one electromyography (EMG) channels. During scoring, an initial artifact rejection step was performed through visual inspection to identify segments requiring exclusion from subsequent analyses.

For subsequent analyses, a subset of 106 EEG channels providing optimal and consistent scalp coverage was selected. These channels underwent automated artifact detection [71], identifying channels exhibiting unusually high or low standard deviations, excessive noise, flat signals, or high-amplitude peaks. Bridged channels were detected based on abnormally high correlations between neighboring channels [72]. Channels were retained only if they were artifact-free for at least 95% of the recording duration across both nights. This procedure resulted in the retention of 89 to 106 channels per participant across nights (mean number of retained channels: RS = 98.52; TMR = 98.43). A final review confirmed comparable scalp coverage across participants.

Spindle events were then automatically detected on each channel using the algorithm described in [14]. To ensure that spindles reflected meaningful neural activity, only events simultaneously detected on 10–40% of the retained channels, with a minimum overlap of 500msec were included. Spindles occurring on fewer than 10% of channels were deemed highly localized events, whereas the ones detected on more than 40% of channels were flagged as potential artifacts. Spindle density was calculated as the number of detected spindles occurring outside auditory stimulation periods ("out" spindles) during N2 and N3 sleep divided by the total duration of these sleep stages. This choice was made to ensure that detected sigma activity reflected genuine spindle events rather than stimulation-related activity.

SOs were identified using the same preprocessing pipeline, with automatic detection [73] performed according to standard parameters outlined in [74]. Only slow oscillations detected on at least 10% of EEG channels were retained for subsequent analyses.

### 2.6. Sleep oscillation analyses

For all analyses, Pearson’s or Spearman’s correlations were performed depending on whether the normality assumption was met.

#### 2.6.1. Spindle density analysis

To allow sufficient time for TMR to influence sleep oscillations, spindle analyses were restricted to N2 and N3 sleep during the second sleep cycle. Spindle density was analyzed using mixed-design ANOVAs, with significant effects followed by Bonferroni-corrected post hoc comparisons.

To investigate the relationship between spindle density and motor performance, correlations were computed between spindle density during the experimental night and the following behavioral measures: (1) GPI at the end of the learning session, calculated as the mean GPI across the final two blocks; (2) mean GPI during the morning retest; and (3) GPI at the beginning of the relearning session, calculated as the mean GPI across the first two blocks. In addition, three relative performance-change indices were calculated. The first quantified the percentage change in performance from the end of learning to the morning retest as: [(mean retest GPI − mean end-of-learning GPI) / mean end-of-learning GPI] × 100. The second quantified the percentage change from the end of learning to the beginning of relearning: [(mean beginning-of-relearning GPI − mean end-of-learning GPI) / mean end-of-learning GPI] × 100. The third quantified the percentage change between the morning retest and the beginning of relearning: [(mean beginning-of-relearning GPI − mean retest GPI) / mean retest GPI] × 100.

To investigate the associations between spindle density and structural plasticity, correlations were computed between spindle density during the experimental night and changes in diffusion MRI metric (see Section 2.8.) between DWI 1 and DWI 2 (pre-learning to post-learning), DWI 1 and DWI 3 (pre-learning to post-night) and between DWI 1 and DWI 4 (pre-learning to delayed retest). Correlations were computed both for predefined bilateral subcortical regions of interest (ROIs), including the cerebellar cortex, thalamus, caudate, putamen, pallidum, and hippocampus, and vertex-wise cortical diffusion metrics. For the cortical analyses, multiple comparisons were corrected using the permutation-based cluster-wise correction procedure described in [62].

#### 2.6.2. Slow oscillation density analysis

SO density was analyzed using an equivalent statistical approach. Because TMR stimulation was initiated during the first sleep cycle, SO density was quantified across N3 sleep from both the first and second sleep cycles. Mixed-design ANOVAs were performed, followed by Bonferroni-corrected post hoc comparisons where appropriate. Correlations between SO density, behavioral measures and diffusion MRI metric changes were conducted using the same approach as described above.

#### 2.6.3. SO–spindle co-occurrence analysis

SO–spindle co-occurrence density was assessed during N3 sleep across the first two sleep cycles. Because TMR stimulation already started during N2 of the first sleep cycle, both cycles were included in the analysis. Co-occurrence density was defined as the proportion of detected spindle events that temporally overlapped with a detected slow oscillation, with analyses restricted to spindles occurring outside auditory stimulation periods. Statistical analyses were performed using the same mixed-design ANOVA framework, followed by Bonferroni-corrected post hoc comparisons where appropriate. Correlations between SO– spindle co-occurrence density, behavioral measures and diffusion MRI metric changes were performed using the same correlation procedures described above.

### 2.7. Sleep Spindle Connectivity Analyses

#### 2.7.1. Source reconstruction and connectivity computation

Source localization was performed using a standard brain template [75] together with a co-registered hd-EEG montage. Source reconstruction was conducted using the Maximum Entropy on the Mean (MEM) framework [76] which is well suited for estimating cortical activity from EEG recordings acquired with variable electrode subsets while remaining non-committal with respect to missing information [77]. The use of a common cortical template (8000 nodes) allowed all participants to be projected onto the same source space, enabling group-level comparisons. The source space was parcellated into 148 cortical regions according to the Destrieux atlas [78]. The inverse problem was solved in the spatial Fourier domain, using a MEM prior defined as a mixture of two zero-mean Gaussian distributions: one with zero variance, corresponding to the inactive state of the Fourier mode, and the other with non-zero variance, corresponding to its active state (see Additional File and LiveScript- localisation available in [79]).

Each detected sleep spindle was divided into two equal-duration segments corresponding to the early and late phases of the spindle. Source activity was estimated independently for each segment, and dimensionality reduction was achieved by averaging the reconstructed source signals within each atlas-defined cortical parcel. For both spindle segments, a functional connectivity graph was then constructed by computing the weighted Phase Lag Index (wPLI) between all pairs of cortical regions. This yielded phase-synchronization networks characterizing the evolution of functional connectivity from the early to the late phase of individual spindle events.

#### 2.7.2. Statistical analysis of spindle connectivity

Statistical analyses were conducted at the group level on individual wPLI connectivity matrices (148-by-148 nodes) or on proportional changes derived from individual wPLI matrices (see below), using the Network-Based Statistic software (NBS; [80, 81]). Statistical contrasts were performed using one-sample, paired-sample, or independent two-sample *t*-tests, as appropriate. NBS is a non-parametric statistical tool that performs multiple univariate tests on all analysed edges (in this case, each element in the connectivity matrix; see [82–84] for similar approaches), while controlling for multiple comparisons across the connectome [80, 81]. Statistical significance of the resulting network components was assessed using 10,000 permutations, with a corrected significance threshold set at pcorr < 0.05.

For each significant network identified through NBS, node-level contributions were quantified using a node strength measure, defined as the sum of all weighted edges connected to a given node [85]. The resulting brain networks and weighted nodes were visualized using the BrainNet Viewer Connectivity Toolbox [86] and the LiveScript ’HD-connectome’ Toolbox [79] that displays Destrieux connectome associated with connectivity metrics. Supplemental one-tailed Pearson’s correlations were computed to investigate associations between mean network connectivity strength and behavioral performance.

##### 2.7.2.1. Within-spindle connectivity dynamics

To characterize the temporal evolution of brain connectivity within spindles in the RS and TMR groups, we computed for each individual wPLI connectivity patterns at the beginning and the end of the out-of-stimulation-periods spindles in sleep stages 2 and 3 (combined) during the second sleep cycle. This provided wPLI connectivity matrices for subsequent network analyses without spatial priors. To identify significant networks in which connectivity evolved from spindle beginning to end, connectivity matrices (148-by-148 nodes) differences were analysed using NBS paired t-tests.

##### 2.7.2.2. Between-night and between-group comparisons of connectivity dynamics

First, we computed for each individual the proportional changes in connectivity patterns from the beginning to the end of the spindles during sleep stages 2 and 3 (combined) during the second sleep cycle. This provided differential connectivity matrices for subsequent network analyses. Statistical contrasts on differential matrices were computed between nights (Night1 vs. Night2 in RS or TMR groups) or groups (RS vs. TMR Sleep in Night1 or Night2). Supplemental one-tailed Pearson’s correlations were computed to investigate associations between mean proportional changes in network connectivity strength and behavioral performance.

##### 2.7.2.3. Correlations with behavioral performance and dMRI metrics

To investigate associations between spindle-related connectivity dynamics and behavioral performance or diffusion MRI changes, we calculated the mean proportional change in connectivity from spindle beginning to end during the experimental night (Night 2). For these analyses, connectivity values were averaged across the edges belonging to the network identified as significant in the NBS contrast of interest. Specifically, because the primary comparison of interest was the difference between the RS and TMR groups during Night 2, the significant edges identified by the between-group unpaired t-test were used as a common network mask. Thus, mean proportional connectivity changes were extracted from the same NBS-derived network for each participant in both groups, allowing direct comparison and correlation with behavioral and diffusion MRI measures.

### 2.8. Diffusion MRI Data Acquisition & Processing

The MRI acquisition parameters and the processing pipeline were overtaken from [62]. dMRI results are published in [62].

## 3. Results

### 3.1. Main behavioral and dMRI findings

All behavioral and diffusion MRI findings were reported in detail in Stee et al. [62]. They are summarized here for the sake of completeness.

At the behavioral level, participants demonstrated robust motor sequence learning during Day 1, with progressive performance improvements and sensitivity to pseudo-random blocks confirming sequence acquisition. While no overnight gains were observed after the experimental night, significant delayed offline improvements emerged after three additional nights, indicating delayed consolidation of the motor memory. During the relearning session on Day 5, participants further improved their performance with practice, reflecting additional online gains. The RS and TMR groups followed comparable performance trajectories throughout the protocol, indicating that TMR did not overtly modulate behavioral performance compared with regular sleep.

At the microstructural level, diffusion MRI revealed dynamic microstructural remodeling associated with motor learning and relearning, characterized by coordinated changes across Diffusion Tensor Imaging (DTI) and Neurite Orientation Dispersion and Density Imaging (NODDI) metrics (see [62]). Initial learning induced widespread decreases in mean diffusivity (MD), accompanied by increases in neurite density index (NDI), decreases in free water fraction (FWF), and region-specific changes in orientation dispersion index (ODI), suggesting increased tissue densification, enhanced neurite density, and local reorganization of fiber architecture. These adaptations were observed across extensive cortical territories and key motor-memory regions, including the putamen, cerebellar cortex, hippocampus, and thalamus. Following the experimental night, several of these learning-related changes persisted, although with reduced magnitude, indicating continued microstructural remodeling during the early consolidation period. Also, some sleep-related modulation emerged, particularly in the right caudate (MD/Fractional Anisotropy (FA)/FWF) and prefrontal cortex (MD/FWF), where RS and TMR groups displayed divergent microstructural trajectories. Three days later, most initial changes had attenuated, suggesting a gradual normalization or refinement of the learning-induced structural response. TMR-related differences were mainly observed in ODI at this stage, particularly in the right cuneus, suggesting possible effects on fiber organization. Relearning elicited a renewed pattern of microstructural adaptations, again characterized by decreased MD, increased NDI, reduced FWF, and ODI alterations, supporting the dynamic and experience-dependent nature of brain plasticity. This effect was comparable between RS and TMR groups, indicating that TMR did not significantly alter relearning-related microstructural remodeling.

### 3.2. Impact of TMR on sleep oscillations & association with dMRI metrics

#### 3.2.1. Spindle density

The mixed-design ANOVA with *Night* (1-Habituation vs. 2-Experimental) as within-subject factor and *Group* (RS vs. TMR) as between-subject factor revealed a main effect of *Night* (*F*_1,57_ = 40.306, *p* < 0.001, η² = 0.414) and *Group* (*F*_1,57_ = 7.198, *p* = 0.010, η² = 0.112), as well as a *Night* × *Group* interaction (*F*_1,57_ = 16.458, *p* < 0.001, η² = 0.224). Post-hoc comparisons showed that spindle density remained stable across nights in the RS group (*p* = 0.646), but markedly decreased in the TMR group (*p* < 0.001). Between-group comparisons further indicated no difference on Night 1 (*p* = 1.000), but significantly lower spindle density in the TMR group on Night 2 as compared to the RS group (*p* < .001; see Figure 3).

**Figure 3:**
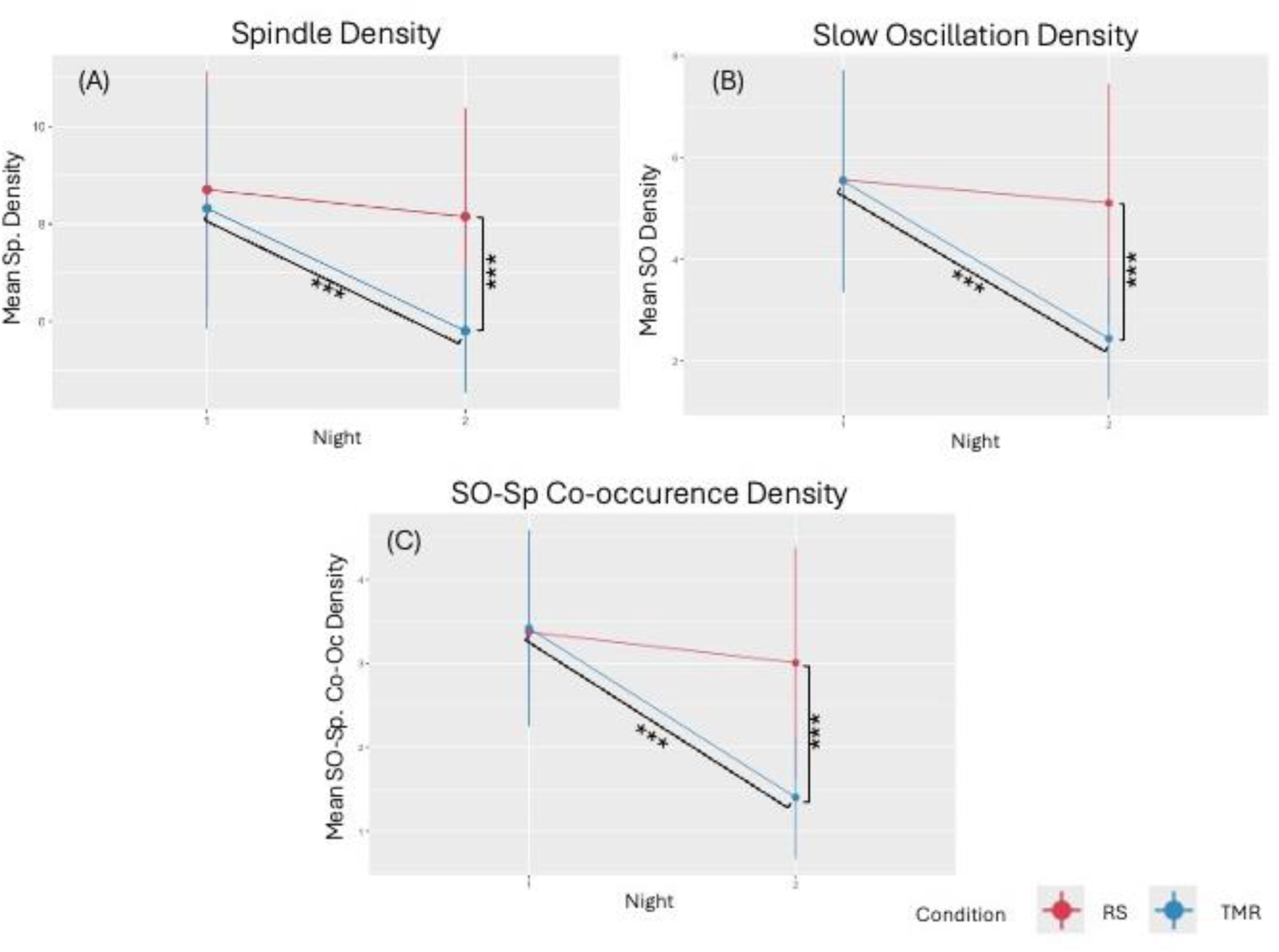
Sleep oscillation densities during the habituation and experimental nights. (A) Spindle density, computed from spindles detected outside auditory stimulation periods, for the Regular Sleep (RS) and Targeted Memory Reactivation (TMR) groups. (B) Slow oscillation (SO) density. (C) SO–spindle co-occurrence density, calculated from spindle events occurring concurrently with a slow oscillation. Data are presented as mean ± sd for the habituation (1) and experimental (2) nights. Significant effects are indicated where applicable (* for *p* < 0.05, ** for *p* < 0.01, *** for *p* < 0.001).

We next examined whether spindle density during the experimental night was associated with behavioral performance. Spindle density positively correlated with the percentage change in performance from the end of the learning session to the morning retest (ρ = 0.384, p = 0.003). No significant associations were observed with the other performance-change indices (all p > 0.072) or with mean GPI values at any of the three time points (all p > 0.141). We next examined whether spindle density during the experimental night correlated with changes in diffusion MRI metrics.

For changes from DWI 1 (pre-learning) to DWI 2 (post-learning), correlations with spindle density were observed across several subcortical ROIs. MD negatively correlated with spindle density in the right caudate (ρ = −0.460, p < 0.001) and right thalamus (ρ = −0.274, p = 0.038). For FA, correlations were observed with the right caudate (ρ = 0.320, p = 0.015), left caudate (ρ = −0.360, p = 0.006), and right thalamus (ρ = 0.389, p = 0.003). ODI positively correlated with spindle density in the left caudate (ρ = 0.436, p < 0.001), right pallidum (ρ = 0.285, p = 0.031), and right putamen (ρ = 0.387, p = 0.003). Finally, FWF negatively correlated with spindle density in the right caudate (ρ = −0.433, p < 0.001) and right thalamus (ρ = −0.260, p = 0.049). No other significant correlations were observed across the subcortical ROIs (all p > 0.057), and no significant clusters emerged in the cortical analysis.

For changes from DWI 1 (pre-learning) to DWI 3 (post-night), correlations with spindle density were observed in subcortical ROIs: MD in the right caudate (ρ = –0.317, *p* = 0.016) and right thalamus (ρ = –0.315, *p* = 0.017); FWF in the right caudate (ρ = –0.375, *p* = 0.004) and left putamen (ρ = 0.264, *p* = 0.048); and ODI in the right putamen (ρ = 0.308, *p* = 0.020). No other significant correlations were found in the ROIs (all *p’*s > 0.084), and no significant clusters emerged in the cortical analysis.

For changes from DWI 1 (pre-learning) to DWI 4 (delayed), spindle density correlated with ODI in the left caudate (ρ = 0.266, *p* = 0.046). No further significant correlations were found in the ROIs (all *p* > 0.123), and no cortical clusters reached significance.

Overall, TMR was associated with a marked reduction in spindle density during the experimental night, whereas spindle density remained stable across nights in the RS group. Higher experimental-night spindle density correlated with greater overnight performance improvement, as well as with multiple learning-related dMRI changes, particularly in the caudate and thalamus. These associations were less extensive at the post-night and delayed time points. No significant associations were detected at the cortical level.

#### 3.2.2. Slow Oscillation density

The mixed-design ANOVA with *Night* (1-Habituation vs. 2-Experimental) as within-subject factor and *Group* (RS vs. TMR) as between-subject factor revealed robust main effects of *Night* (*F*_1,57_ = 38.846, *p* < 0.001, η² = 0.405) and *Group* (*F*_1,57_ = 13.017, *p* < 0.001, η² = 0.186), together with a *Night* × *Group* interaction (*F*1,57 = 21.789, *p* < 0.001, η² = 0.277). Post-hoc tests indicated that SO density was unchanged across nights in the RS group (*p* = 1.000), but showed a clear reduction from Night 1 to Night 2 in the TMR group (*p* < 0.001). Moreover, while the RS and TMR groups did not differ at baseline (Night 1, *p* = 1.000), the TMR group exhibited lower SO density than the RS group during the experimental night (Night 2, *p* < .001; see Figure 3).

We next examined whether SO density during the experimental night correlated with behavioral performance. No significant correlations were observed between SO density and either raw GPI values or any of the performance-change indices (all *p*’s > 0.076).

We then investigated whether SO density during the experimental night was associated with changes in diffusion MRI metrics across the different post-learning timepoints.

For changes from DWI 1 (pre-learning) to DWI 2 (post-learning), correlations with SO density were observed across several subcortical ROIs. MD negatively correlated with SO density in the left cerebellar cortex (r = −0.341, p = 0.009), left hippocampus (r = −0.450, p < 0.001), right caudate (ρ = −0.397, p = 0.002), right cerebellar cortex (ρ = −0.264, p = 0.046), and right thalamus (ρ = −0.350, p = 0.007). For FA, correlations were observed in the left (ρ = −0.346, p = 0.009) and right caudate (ρ = 0.301, p = 0.022). ODI positively correlated with SO density in the left caudate (ρ = 0.316, p = 0.017) and right pallidum (r = 0.268, p = 0.042). Finally, FWF negatively correlated with SO density in the left hippocampus (r = −0.263, p = 0.046), right caudate (ρ = −0.386, p = 0.003), and right thalamus (ρ = −0.375, p = 0.004). No additional significant correlations were observed across the subcortical ROIs (all *p’*s > 0.057), and no significant clusters emerged in the cortex.

For changes from DWI 1 (pre-learning) to DWI 3 (post-night), several correlations emerged in subcortical ROIs: with FWF in the left caudate (ρ = 0.302, *p* = 0.023); with MD in the right caudate (ρ = –0.281, *p* = 0.033) and right thalamus (r = –0.258, *p* = 0.050); and with ODI in the right pallidum (ρ = 0.288, *p* = 0.029). No additional ROI correlations reached significance (all *p*’s > 0.062). In the cortex, there was a positive correlation between ODI and SO density in a cluster located in the right fronto-marginal gyrus & sulcus.

For changes from DWI 1 (pre-learning) to DWI 4 (delayed), SO density positively correlated with FA in the left pallidum (ρ = 0.316, *p* = 0.016). No further significant associations were found in the ROIs (all *p* > .071), and no cortical clusters reached significance.

To sum up, TMR was associated with a marked reduction in SO density during the experimental night, whereas SO density remained stable across nights in the RS group. Higher experimental-night SO density was associated with multiple learning-related dMRI changes, particularly in the caudate, thalamus, hippocampus, and cerebellar cortex. These associations were most extensive at the immediate post-learning time point and became substantially more limited at the post-night and delayed time points, with only isolated subcortical and cortical associations persisting. Despite these associations with microstructural changes, SO density was not directly related to behavioral performance or its evolution across the experimental period.

#### 3.2.3. SO-spindle co–occurrence density

The mixed-design ANOVA with *Night* (1-Habituation vs. 2-Experimental) as within-subject factor and *Group* (RS vs. TMR) as between-subject factor revealed main effects of *Night* (*F*_1,57_ = 46.690, *p* < 0.001, η² = 0.450), *Group* (*F*_1,57_ = 13.889, *p* < 0.001, η² = 0.196), as well as a *Night* × *Group* interaction (*F*_1,57_ = 22.837, *p* < 0.001, η² = 0.286). Post-hoc tests showed that SO– spindle co-occurrence density remained stable from Night 1 to Night 2 in the RS group (*p* = 0.890), but markedly decreased from Night 1 to Night 2 in the TMR group (*p* < 0.001). Importantly, the two groups did not differ on Night 1 (*p* = 1.000), whereas the TMR group exhibited lower density than the RS group on Night 2 (*p* < 0.001; see Figure 3).

We next examined whether SO–spindle co-occurrence density during the experimental night correlated with behavioral performance. Co-occurrence density positively correlated with the percentage change in performance from the end of the learning session to the morning retest (ρ = 0.257, p = 0.050). No significant associations were observed with the other performance- change indices (all p > 0.209) or with raw GPI values at any of the assessed time points (all p > 0.601).

We then investigated whether SO–spindle co-occurrence density during the experimental night was associated with changes in diffusion MRI metrics across the different post-learning time points.

For changes from DWI 1 (pre-learning) to DWI 2 (post-learning), correlations with SO–spindle co-occurrence density were observed across several subcortical ROIs. MD negatively correlated with SO–spindle co-occurrence density in the left (ρ = −0.314, p = 0.017) and right cerebellar cortex (ρ = −0.272, p = 0.039), left hippocampus (ρ = −0.354, p = 0.007), right caudate (ρ = −0.393, p = 0.002), and right thalamus (ρ = −0.340, p = 0.009). For FA, correlations were observed in the left (ρ = −0.354, p = 0.007) and right caudate (ρ = 0.271, p = 0.040). ODI positively correlated with SO–spindle co-occurrence density in the left caudate (ρ = 0.321, p = 0.015) and right putamen (ρ = 0.295, p = 0.027). FWF was negatively correlated with SO– spindle co-occurrence density in the right caudate (ρ = −0.383, p = 0.003) and right thalamus (ρ = −0.361, p = 0.006). No other significant correlations were observed across the subcortical ROIs (all p > 0.120), and no significant clusters were identified in the cortical analysis.

For changes from DWI 1 (pre-learning) to DWI 3 (post-night), a correlation was observed in the right caudate for FWF (ρ = –0.297, *p* = 0.024). No other ROI correlations reached significance (all *p* > 0.053), and no significant clusters were identified in the cortical analysis.

For changes from DWI 1 (pre-learning) to DWI 4 (delayed), SO–spindle co-occurrence density correlated with FA in the left pallidum (ρ = 0.265, *p* = 0.045). No further significant correlations were observed in the ROIs (all *p* > .070), and no significant cortical clusters emerged.

Hence, TMR markedly reduced SO–spindle co-occurrence density during the experimental night, whereas co-occurrence remained stable across nights in the RS group. Higher co-occurrence density was associated with greater overnight performance improvement, although this association was modest and was not observed for the other behavioral measures. Co-occurrence density was also associated with multiple learning-related dMRI changes immediately after learning, particularly in the cerebellar cortex, hippocampus, caudate, and thalamus. These associations became substantially less extensive at the post-night and delayed time points, with only isolated subcortical associations persisting and no significant cortical associations.

#### 3.2.4. SO-spindle co-occurrence & spindle frequency

The mixed-design ANOVA with *Night* (1-Habituation vs. 2-Experimental) and *Co-occurrence* (Yes vs. No) as within-subject factors, and *Group* (RS vs. TMR) as between-subject factor, performed on spindle frequency during N3 across both cycles, revealed a main effect of *Night* (*F*_1,57_ = 4.195, *p* = 0.045, η² = 0.069). Post-hoc comparisons indicated a slower spindle frequency on the first night (mean = 12.722 Hz, SD = 0.300) compared to the second night (mean = 12.758 Hz, SD = 0.316). All other main effects (all *p* > 0.314) and interactions (all *p* > 0.142) were not significant.

Overall, spindle frequency showed only a restricted but significant increase from the habituation to the experimental night, with no evidence that TMR or SO–spindle co-occurrence modulated spindle frequency.

3.3. Evolution of cortico-cortical connectivity during sleep spindles and correlation with dMRI;

#### 3.3.1. Temporal evolution of brain connectivity within spindles

Using NBS [80, 81] paired *t*-tests revealed significantly greater wPLI connectivity during the end as compared to the beginning of sleep spindles during the baseline night (Night 1) in both the RS and TMR groups, consistent with previous reports [55]. In the RS group, two significant subnetworks were identified, whereas one significant subnetwork emerged in the TMR group. Across these subnetworks, the strongest nodes were distributed across fronto-central, sensorimotor, cingulate, insular, parietal, temporal, and occipital cortices, including the precentral and postcentral gyri, orbitofrontal and superior frontal regions, anterior and posterior cingulate cortex, insula, precuneus, superior and inferior parietal cortices, superior temporal cortex, and visual occipital areas.

During the experimental night (Night 2), the RS group continued to exhibit increased connectivity from the beginning to the end of the spindles. The resulting network involved widespread sensorimotor, occipital, parietal, temporal, orbitofrontal, and cingulate regions, with the strongest node contributions arising from the precentral gyrus, superior and middle occipital cortex, orbitofrontal cortex, superior temporal cortex, angular gyrus, postcentral gyrus, and precuneus. At variance, no significant differences in connectivity between the early and late phases of the spindle were observed in the TMR group during the experimental night. The complete lists of nodes, node strength values, and Montreal Neurological Institute (MNI) coordinates are provided in the Additional File (Additional Tables 1–3 and Additional Figures 1–3).

To summarize, intra-spindle connectivity increased from the beginning to the end of the spindles across widespread cortical networks during the baseline night in both groups and persisted during the experimental night in the RS group. In contrast, this connectivity evolution was no longer detectable during the experimental night in the TMR group, suggesting that TMR altered the normal temporal dynamics of cortical connectivity across the spindle.

#### 3.3.2. Between-night changes in the evolution of brain connectivity within spindles

Using NBS [80, 81], paired *t*-tests comparing the proportional change in wPLI connectivity between the beginning and the end of the spindles across nights revealed opposite patterns in the two groups. In the RS group, proportional increases in within-spindle connectivity were significantly greater during the experimental night (Night 2) as compared with the baseline night (Night 1). This network predominantly involved fronto-insular, cingulate, parietal, and occipital regions, with prominent contributions from the anterior insula, inferior parietal/angular gyri, posterior cingulate cortex, superior occipital cortex, orbitofrontal regions, and sensorimotor areas.

In contrast, the TMR group exhibited greater proportional increases in within-spindle connectivity during the baseline night compared with the experimental night, reflecting a reduction in the normal spindle-related connectivity evolution following TMR. This network was more extensive and included widespread fronto-temporal, parietal, insular, cingulate, sensorimotor, and occipital regions, with strong contributions from the posterior cingulate cortex, inferior temporal cortex, frontal pole/transverse frontopolar regions, superior temporal cortex, supramarginal and postcentral regions, insula, and orbitofrontal cortex. The complete lists of nodes, node strength values, and MNI coordinates are provided in the Additional File (Additional Tables 4–5 and Additional Figures 4-5).

**Figure 4.**
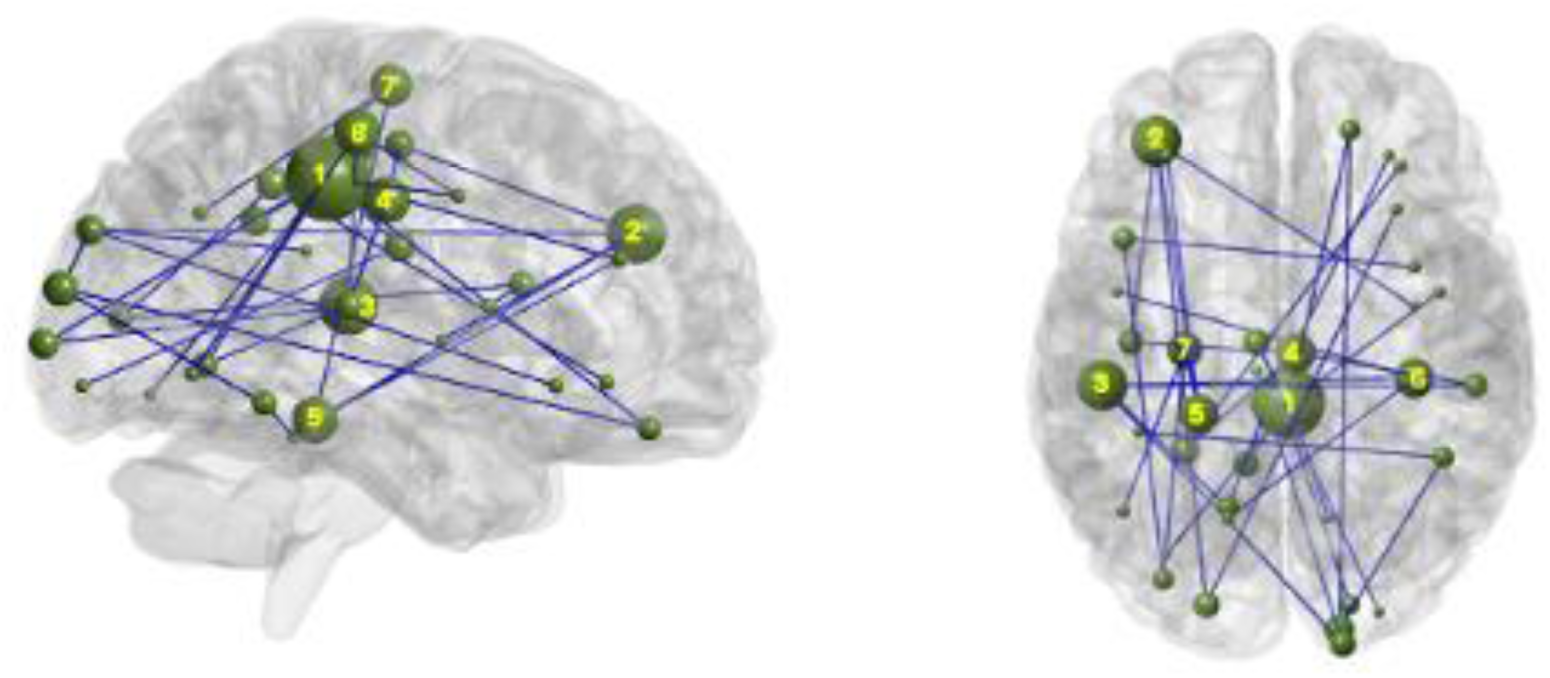
Between-group differences in proportional connectivity changes from spindle beginning to end. Node size represents node strength differences between groups (RS > TMR). Only the seven nodes with the highest strength are shown for legibility, see Additional Table 6 for a complete list with coordinates. Displayed location labels [78] 1: G_cingul-Post-dorsalR; 2: G_front_middleL; 3: S_temporal_transverseL; 4: G_and_S_cingul-Mid-PostR; 5: G_oc-temp_med-ParahipL; 6: G_postcentralR ; 7: S_precentral-sup-partL

Overall, the two groups exhibited opposite night-to-night changes in intra-spindle connectivity evolution: connectivity increases from spindle onset to termination became more pronounced during the experimental night in the RS group, but were reduced in the TMR group. These findings further indicate that TMR disrupted the normal night-to-night enhancement of connectivity dynamics within sleep spindles.

#### 3.3.3. Between group changes in the evolution of brain connectivity within spindles

During the baseline night (Night 1), no significant differences in proportional connectivity changes between the beginning and the end of the spindles were observed between the RS and TMR groups. During the experimental night (Night 2), however, the RS group exhibited significantly greater spindle-related connectivity evolution compared with the TMR group. The identified network comprised widespread cingulo-frontal, sensorimotor, temporal, parietal, and occipital regions, with prominent contributions from the posterior cingulate cortex, middle frontal gyrus, temporal cortex, parahippocampal region, postcentral and precentral gyri, superior occipital cortex, angular/inferior parietal cortex, insular cortex, and orbitofrontal areas (Figure 4). The complete lists of nodes, node strength values, and MNI coordinates are provided in the Additional File (Additional Table 6 and Additional Figure 6).

Overall, the two groups did not differ in spindle-related connectivity dynamics at baseline, but TMR was associated with a marked reduction in connectivity evolution during the experimental night, as compared to greater spindle-related connectivity changes in the RS group across a widespread cortical network.

#### 3.3.4. Correlation of intra-spindle connectivity evolution with performance

We next examined whether intra-spindle connectivity evolution during the experimental night was associated with behavioral performance. No significant correlations were observed between intra-spindle connectivity evolution and GPI values measured at the end of the learning session (Day 1), during the morning retest (Day 2), or at the beginning of the relearning session (Day 5; all p > 0.154). Similarly, no significant associations were found between intra-spindle connectivity evolution and relative changes in performance between the morning retest and the end of the learning session or between the delayed retest and the end of the learning session (all p > 0.120).

Overall, intra-spindle connectivity evolution during the experimental night was not significantly associated with either motor performance or changes in performance across the post-learning time points.

#### 3.3.5. Correlation of intra-spindle connectivity with dMRI metrics

Correlations were performed between proportional changes in wPLI connectivity from the beginning to the end of sleep spindles during the experimental night (Night 2) and diffusion MRI changes following learning (DWI 2 – DWI 1). Significant correlations were observed in several subcortical ROIs. In the left caudate, intra-spindle connectivity changes correlated negatively with FA changes (ρ = –0.291, *p* = 0.027) and positively with ODI changes (ρ = 0.383, *p* = 0.003). In the left hippocampus, connectivity changes negatively correlated with MD changes (ρ = –0.364, *p* = 0.005), while in the left putamen, correlations were observed with MD (*r* = –0.265, *p* = 0.049) and NDI (*r* = 0.319, *p* = 0.017). In the right caudate, connectivity changes negatively correlated with MD (ρ = –0.406, *p* = 0.001) and FWF (ρ = –0.349, *p* = 0.007), and positively with FA (*r* = 0.325, *p* = 0.012). Finally, a positive correlation was observed between intra-spindle connectivity changes and FA changes in the right thalamus (*r* = 0.311, *p* = 0.016). No additional subcortical ROI correlations reached significance (all *p* > 0.051) and no significant correlations were identified in cortical surface analyses.

Correlations were also performed between proportional changes in wPLI connectivity from the beginning to the end of sleep spindles during the experimental night (Night 2) and post-night diffusion MRI changes (DWI 3 – DWI 1). Significant associations were observed in several subcortical regions. In the left thalamus, intra-spindle connectivity changes correlated positively with FA (*r* = 0.298, *p* = 0.022) and NDI (*r* = 0.268, *p* = 0.040). In the right caudate, connectivity changes negatively correlated with MD (*r* = –0.317, *p* = 0.014) and FWF (*r* = – 0.300, *p* = 0.021). In the right putamen, negative correlations were observed with MD (ρ = – 0.262, *p* = 0.047), while ODI showed a positive correlation (*r* = 0.300, *p* = 0.022). Finally, in the right thalamus, intra-spindle connectivity changes negatively correlated with MD (*r* = –0.319, *p* = 0.014) and positively with NDI (ρ = 0.336, *p* = 0.009). No additional subcortical ROI correlations reached significance (all *p* > 0.053). Cortical surface analysis revealed one positively significant cluster in the right pericallosal sulcus for ODI changes. No significant cortical clusters were identified for FA, MD, NDI, left hemisphere ODI, or FWF.

Correlations were performed between proportional changes in wPLI connectivity from the beginning to the end of sleep spindles during the experimental night (Night 2) and delayed diffusion MRI changes (DWI 4 – DWI 1). Significant correlations were observed in several subcortical ROIs. In the left caudate, intra-spindle connectivity changes correlated positively with ODI changes (ρ = 0.294, *p* = 0.025). In the left cerebellar cortex, connectivity changes negatively correlated with FWF changes (*r* = –0.256, *p* = 0.050). Similarly, a negative correlation was observed between intra-spindle connectivity changes and FWF changes in the right caudate (*r* = –0.268, *p* = 0.040). No additional significant correlations were identified across the remaining subcortical ROIs (all *p* > 0.063), and no significant cortical clusters emerged.

Overall, greater intra-spindle connectivity evolution was associated with multiple subcortical dMRI changes following learning, particularly in the caudate, thalamus, hippocampus, and putamen. Several associations persisted at the post-night time point, including a significant cortical association in the right pericallosal sulcus, whereas only a small number of subcortical associations remained at the delayed time point.

## 4. Discussion

The primary objective of this study was to investigate the effects of TMR on sleep oscillations and their associations with memory consolidation and dMRI-derived measures of brain microstructure. We examined spindle density, SO density, SO–spindle co-occurrence, and spindle frequency. Our results show a marked reduction in spindle density, SO density, and SO–spindle co-occurrence during the experimental night in the TMR group, whereas these measures remained stable across nights in the RS group. Spindle frequency moderately increased across nights that were not modulated by TMR. Higher spindle density and SO– spindle co-occurrence during the experimental night were associated with greater improvement in performance from the end of learning to the morning retest, whereas SO density alone was not. All three sleep oscillatory measures had multiple associations with learning-related dMRI changes, particularly between the pre-learning (DWI 1) and post-learning (DWI 2) scans. These associations were most prominent in subcortical regions, including the caudate, thalamus, hippocampus, putamen, pallidum, and cerebellar cortex, and became less extensive at the post-night (DWI 3) and delayed (DWI 4) time points. Few and no consistent associations were observed at the cortical level.

The second objective was to characterize the temporal evolution of cortico-cortical connectivity across individual sleep spindles, and determine whether it was modulated by learning and TMR. During the baseline night, both groups showed a widespread increase in cortico-cortical connectivity from the beginning to the end of individual spindles. Following learning, this intra-spindle connectivity evolution was enhanced in the RS group but attenuated following TMR. Direct between-group comparisons confirmed greater connectivity evolution in the RS than TMR group during the experimental night, with no group difference at baseline. Although intra-spindle connectivity evolution was not directly associated with behavioral performance, it had multiple associations with learning-related dMRI changes, particularly in subcortical regions. Overall, these findings suggest that our experimental TMR protocol did not simply fail to enhance the physiological signatures of memory consolidation, but instead altered several aspects of sleep microstructure and spindle-related network dynamics during the post-learning night.

### 4.1. TMR-induced alterations in sleep oscillatory dynamics

The reduction in spindle density, SO density, and SO–spindle co-occurrence following TMR contrasts with the more commonly reported enhancement of sleep oscillations following successful auditory reactivation. One possible explanation is that the efficacy of TMR depends critically on the temporal relationship between cue presentation and endogenous sleep dynamics. In particular, successful memory reactivation has been linked to the precise timing of auditory cues relative to ongoing spindle activity [39, 87, 88]. Mistimed or too frequent stimulation may therefore not simply render TMR ineffective, but actively interfere with the sleep processes initiated by previous cues.

Several studies support this possibility. Closely spaced auditory cues have been shown to interrupt the emergence or continuation of sleep spindles and to prevent associated memory benefits [38, 89]. In particular, auditory cues separated by at least 1300 ms enhanced sigma activity, whereas shorter inter-cue intervals reduced sigma-band activity. Comparable effects have been reported in rodents, where presentation of a subsequent cue interrupted either memory reactivation [30] or an ongoing spindle [90]. Together, these findings suggest that successive cues need to be sufficiently separated to allow the neurophysiological processes initiated by one cue to unfold before another stimulus is delivered. Our stimulation protocol may therefore have been suboptimal for preserving endogenous spindle dynamics. We attempted to reactivate the entire learned sequence using a 6-sec auditory sequence composed of 12 consecutive 500-ms cues. Although this approach was intended to maximize reactivation of the complete motor sequence, with short interstimuli intervals aimed at preserving the perceived continuity and structure of the sequence, the resulting relatively prolonged and temporally dense stimulation may instead have interfered with the emergence or maintenance of sleep spindles. This interpretation is consistent with the marked reduction in spindle density observed in the TMR group during the experimental night. Thus, rather than enhancing spindle occurrence, repeated stimulation within a short time window may have disrupted the spindle dynamics required for effective memory processing.

The timing of stimulation relative to endogenous spindle activity may provide an additional explanation. Antony et al. [39] reported that cues delivered less than 3 s after the end of a spindle failed to produce beneficial effects, potentially reflecting a refractory period of the thalamocortical networks involved in spindle generation. Conversely, cues delivered more than 6 sec after a preceding spindle were less likely to elicit a spindle, suggesting that the system may progressively enter a less spindle-permissive state. A window of approximately 3–6 sec after spindle termination therefore appeared to provide the most favorable conditions for spindle enhancement and memory consolidation. Our 6-sec-on/6-sec-off stimulation protocol may consequently have repeatedly presented cues outside this optimal window. Because stimulation was delivered continuously for 6 sec followed by 6 sec of silence, individual cues could have occurred during periods in which the thalamocortical system was either refractory or no longer optimally responsive to spindle induction. Moreover, the stimulation period itself may have overlapped with naturally occurring spindles, potentially interfering with their development or continuation. Thus, both the duration and temporal structure of the stimulation may have contributed to the reduction in spindle density observed here.

A similar consideration may apply to the reduction in SO density. Auditory stimulation during sleep has been shown to alter slow-wave activity and slow-wave propagation under some conditions, with reductions in slow-wave occurrence and spatial propagation being associated with impaired memory performance [91]. The phase of the SO at which stimulation is delivered appears to be particularly important. Beneficial effects have most consistently been reported when cues are presented during the SO upstate, when cortical excitability is relatively high [27, 92–96], whereas findings regarding stimulation during the SO downstate are more heterogeneous, with reports of beneficial [40], detrimental [92], or null effects [97, 98].

Importantly, the effect of stimulation may depend not only on SO phase but also on its relationship with spindle activity. Baxter et al. [28], for example, reported memory benefits when cues were delivered in temporal alignment with SO–spindle coupling, close to the SO upstate peak and during periods of relatively low sigma power. Consistent with this, stimulation during the SO upstate has been associated with increases in delta power [27, 93, 94], sigma activity [92, 94, 98] and SO–spindle coupling [27, 95, 96]. In the present study, however, stimulation was delivered without real-time detection of either SO phase or spindle activity. Cues may therefore have occurred at suboptimal phases of the SO, during periods unfavorable for spindle generation, or outside the temporal window supporting SO–spindle coordination. Rather than reinforcing endogenous slow–fast oscillatory coupling, stimulation may consequently have disrupted it. At the same time, presenting stimuli at about 2 seconds apart to match SO upstates, may have limited the extent to which the sequential structure of the material could have been detected during sleep. Indeed, previous work suggests that the maintenance of external inputs over short periods of time may be altered during sleep [99], preventing the integration of individual cues into more complex sequential representations.

### 4.2. Association with motor memory consolidation

In the motor domain, sleep-dependent gains in performance have repeatedly been associated with sleep spindle density [6, 100–102]. This association is further supported by a recent meta-analysis showing a significant relationship between spindle activity and procedural memory performance (r = 0.319) [103]. Beyond providing a favorable neurophysiological environment for systems consolidation, spindles may also carry information related to the memories being reactivated and processed during sleep [103], potentially by supporting the local reactivation and functional integration of distributed but interconnected regions involved in motor memory, including the hippocampus, striatum, thalamus, and motor-related cortical areas [7]. Similarly, SO–spindle coupling has been consistently associated with sleep-dependent memory gains [20, 22–24, 28, 104], supporting the importance of coordinated slow oscillation–spindle dynamics for memory consolidation [105]. Consistent with this literature, higher spindle density and greater SO–spindle co-occurrence during the experimental night were associated with greater improvement in performance from the end of learning on Day 1 to the morning retest on Day 2 (spindle density: ρ = 0.384, p = 0.003; SO– spindle co-occurrence: ρ = 0.257, p = 0.050). In contrast, SO density alone was not associated with overnight performance changes, consistent with the more variable evidence linking SO occurrence per se to memory consolidation. These findings therefore reinforce the possibility that the coordinated occurrence of spindles and SOs, rather than the expression of either oscillation in isolation, is particularly relevant for motor memory consolidation.

Interestingly, none of the sleep oscillatory measures was associated with raw behavioral GPI scores at any time point. This dissociation may reflect the fact that absolute performance and overnight changes in performance capture different aspects of memory processing.

Associations between performance at the end of training and subsequent sleep oscillations could reflect the strength or amount of information available for reactivation during sleep. Indeed, weaker memories may receive greater prioritization during subsequent sleep, whereas stronger memories may show smaller subsequent gains because of reduced room for improvement or ceiling effects [11, 106]. In contrast, associations between sleep oscillations and overnight changes in performance may more directly reflect the processes involved in post-learning consolidation. Thus, the relationship observed here with overnight performance gains, but not absolute performance, may be more consistent with a role of spindle-related activity in the consolidation process itself.

Overall, our findings provide further evidence that spindle-related activity, particularly when temporally coordinated with SOs, is associated with subsequent motor memory consolidation.

### 4.3. Associations between sleep oscillations and learning-related microstructural changes

Evidence suggests that sleep oscillations are not only a reflection of the functional dynamics of neural networks, but are also influenced and constrained by the structural properties of the underlying brain regions [47–50]. In particular, thalamo-cortical white-matter microstructure [48] and hippocampal gray-matter volume have been positively associated with sleep spindle activity [49, 52], whereas gray-matter volume in the hypothalamus and medial prefrontal cortex has been related to SO activity [52].

In line with these observations, we found that learning-related microstructural changes between DWI 1 and DWI 2 were associated with the expression of sleep oscillations during the subsequent experimental night. Greater decreases in MD and FWF in the caudate and thalamus were associated with higher spindle, SO, and SO–spindle co-occurrence densities. SO density and SO–spindle co-occurrence were additionally associated with MD and FWF changes in the cerebellar cortex and hippocampus, whereas these associations were not observed for spindle density alone. Changes in FA and ODI in the caudate were also associated with sleep oscillatory density, suggesting that the relationship between learning-related microstructural remodeling and subsequent sleep oscillations extends beyond changes in tissue density (reflected by MD & FWF) to alterations in the local integrity and dispersion of fibers (reflected by FA & ODI).

Both the temporal and anatomical profiles of these associations are particularly informative. Temporally, the largest number of associations was observed for microstructural changes occurring during learning itself (DWI 2 – DWI 1), before the experimental sleep episode. These associations therefore cannot represent structural consequences of the subsequent sleep oscillations and instead suggest that learning-induced remodeling may contribute to shaping the neural substrate on which subsequent oscillatory activity operates. Correlations became considerably less extensive for post-night changes (DWI 3–DWI 1), with MD and FWF associations mainly retained in the caudate, thalamus, and putamen, and were largely absent at the delayed time point (DWI 4–DWI 1).

The anatomical distribution of these associations can likewise be relevant to the proposed reorganization of motor memories during sleep. At the immediate post-learning time point (DWI 2 – DWI 1), hippocampal microstructural changes were associated with SO and SO– spindle densities, whereas the caudate and thalamus showed associations across all oscillatory measures. Following the experimental night (DWI 3 – DWI 1), associations were largely restricted to the caudate and thalamus, with no remaining associations involving the hippocampus. Although this pattern should be interpreted cautiously, it is compatible with a progressive shift from an initially more hippocampally dependent representation toward a more distributed subcortico-cortical network during motor memory consolidation [107, 108]. In this framework, learning-related changes in the hippocampus may be associated with the oscillatory activity supporting the initial processing of the newly encoded memory, whereas later consolidation may increasingly involve subcortical structures such as the caudate and thalamus.

Such interpretation is consistent with the proposed role of sleep spindles in promoting neuronal plasticity. Spindle activity has been implicated in long-lasting changes in neuronal responsiveness and in cellular processes underlying synaptic plasticity (for review, see [109]). Moreover, spindles occurring in conjunction with SO up-states have been associated with a distinct pattern of calcium activity compared with isolated spindles or SOs, potentially providing a particularly favorable state for dendritic plasticity [110]. In the context of motor memory, Boutin et al. [7] proposed that the shift from hippocampal to subcortico-cortical involvement during spindle–ripple events may reflect the concurrent consolidation and redistribution of motor memory representations, alongside the progressive clearance or downscaling of the hippocampal trace. Such reorganization could help prevent synaptic saturation while preserving the capacity to encode new information during subsequent wakefulness, consistent with the broader framework of memory-specific synaptic strengthening occurring alongside global synaptic downscaling during sleep (for review, see [111]).

Taken together, the strongest associations were observed between microstructural changes established during learning and the sleep oscillations expressed during the subsequent night, whereas more limited associations persisted into the post-night period. This pattern suggests a close temporal relationship between learning-induced microstructural remodeling and subsequent sleep oscillatory activity, potentially followed by further microstructural reorganization during sleep-dependent consolidation. However, the correlational nature of these analyses prevents determining whether learning-induced microstructural changes influence subsequent oscillatory activity, whether sleep oscillations contribute to later microstructural remodeling, or whether both reflect a common underlying plasticity process. Nevertheless, the temporal and anatomical distribution of the associations points to a dynamic interaction between learning-related microstructural plasticity and sleep oscillatory activity, particularly within subcortical structures implicated in motor memory consolidation.

### 4.4. Spindle frequency as a function of SO-spindle coupling

Several studies reported associations between fast spindle activity and memory consolidation [101, 112]. However, a recent meta-analysis found no evidence that spindle type (fast vs. slow) systematically moderates the strength of the spindle–memory association on its own, suggesting that both spindle types may contribute to memory consolidation [103]. Nevertheless, Muehlroth et al. [23] showed that, in young adults, memory consolidation was associated with the precise coupling of fast spindles (12.5–16 Hz) to the SO up-state, whereas increased slow spindle activity (9–12.5 Hz) toward the end of the SO up-state was associated with poorer memory consolidation, a pattern more characteristic of older adults.

Given these mixed findings, we conducted an exploratory analysis to determine whether spindles occurring in conjunction with SOs differed in frequency from spindles occurring independently of SOs. We found no significant difference in spindle frequency as a function of SO–spindle co-occurrence, with no evidence that coupled spindles were systematically faster or slower than uncoupled spindles. Thus, although the temporal coupling of spindles with SOs was associated with motor performance, this coupling was not accompanied by a detectable shift in spindle frequency in the present study. This suggests that the functional relevance of SO–spindle coupling may depend more strongly on when spindles occur relative to the SO cycle than on a systematic change in their intrinsic frequency.

### 4.5. Intra-spindle connectivity dynamics

Zerouali et al. [55] showed that connectivity evolves across the course of a sleep spindle, with earlier spindle phases predominantly characterized by short-range, intra-hemispheric connectivity, whereas later phases are associated with stronger and more widespread long-range connectivity. In the present study, we similarly examined the temporal evolution of connectivity across individual spindles. Rather than characterizing changes in the spatial organization of the network, we quantified changes in connectivity strength between the beginning and end of each spindle. Consistent with Zerouali et al.’s report [55], both groups showed an increase in connectivity strength from the early to the late spindle phase during the baseline night, indicating a progressive strengthening of cortico-cortical interactions across spindles.

Following motor sequence learning, this intra-spindle connectivity evolution was further enhanced in the RS group. Connectivity increased more strongly from the early to the late spindle phase during the experimental night than during baseline, suggesting that recent learning modulated the temporal dynamics of spindle-related cortical communication. This finding is compatible with the proposed role of spindles in coordinating the reactivation and functional integration of distributed brain regions involved in motor memory, including cortical motor regions, the striatum, thalamus, and hippocampus [3, 4, 6, 7]. Our findings extend this literature by suggesting that learning may influence not only the occurrence or magnitude of spindle activity, but also its temporal evolution within individual spindle events.

To the best of our knowledge, the effect of TMR on this temporal evolution has not previously been investigated. In contrast to the learning-related increase observed in the RS group, the TMR group showed a reduction in intra-spindle connectivity evolution from baseline to the experimental night. Direct between-group comparisons confirmed greater connectivity evolution in the RS than TMR group during the experimental night, despite no group difference at baseline. Thus, the effects of TMR were not limited to a reduction in spindle occurrence, but extended to the temporal organization of cortical connectivity during individual spindles. This finding suggests that acoustic stimulation may alter aspects of spindle dynamics beyond their frequency of occurrence, potentially interfering with the progressive coordination of cortical activity normally observed across spindle events.

The cortical networks contributing to this evolution also differed between conditions. In the RS group, the identified network was relatively focused, with prominent contributions from the anterior insula, angular/inferior parietal cortex, posterior cingulate cortex, occipital cortex, and orbitofrontal regions. In contrast, the TMR-related network was more spatially extensive, encompassing temporal and occipital cortices, the insula, frontal regions, and posterior cingulate and parietal areas. Because our analysis specifically quantified the evolution of connectivity strength rather than the spatial organization of spindle networks per se, these differences should be interpreted as differences in the cortical regions contributing to connectivity evolution rather than as evidence that TMR recruited an entirely distinct spindle network.

The present findings can be compared with those of Boutin et al. [6], although methodological differences limit direct comparison. Boutin et al. assessed spindle-band functional connectivity within predefined cortical and subcortical ROIs, including primary motor and somatosensory cortices, thalamus, putamen, hippocampus, caudate nucleus, and cerebellum. They reported increased spindle-band coherence within bilateral subcortical networks, together with unilateral interactions between subcortical and sensorimotor cortical regions. In contrast, our analysis examined connectivity across the entire cortical surface and specifically quantified its evolution within individual spindles. Thus, while both studies support the involvement of distributed networks in spindle-related processing following motor sequence learning, they capture different aspects of spindle communication: connectivity strength between predefined cortical and subcortical regions across spindle events in Boutin et al., versus the temporal evolution of cortico-cortical connectivity within individual spindles in the present study.

Consistent with this distinction, we did not observe a direct association between the magnitude of intra-spindle connectivity evolution and motor performance. Boutin et al. [6], in contrast, found that spindle-related cortico-subcortical connectivity was associated with the degree of overnight motor sequence consolidation. This discrepancy may indicate that the temporal evolution of cortico-cortical connectivity within individual spindles is not, in itself, a direct determinant of behavioral consolidation, but may instead represent one component of a broader network-level process supporting memory reorganization during Non-Rapid Eye Movement sleep (NREM).

Interestingly, greater intra-spindle connectivity evolution was associated with multiple learning-related microstructural changes across subcortical regions. Immediately following learning (DWI 2–DWI 1), associations were observed in the caudate, hippocampus, putamen, and thalamus, involving several diffusion metrics that capture complementary aspects of tissue microstructure and organization. Several of these associations persisted following the experimental night (DWI 3–DWI 1), particularly in the caudate, putamen, and thalamus, whereas hippocampal associations were no longer observed. This pattern may reflect an interaction between the structural changes established during learning and the functional network reorganization occurring during subsequent sleep. In particular, the disappearance of hippocampal associations alongside the persistence of associations in the caudate and thalamus is compatible with the broader proposal of a progressive transition from initially hippocampally influenced representations toward more distributed subcortico-cortical networks during motor memory consolidation [6, 107]. However, this interpretation remains speculative, considering the correlational nature of our analyses.

Taken together, these findings suggest that motor learning enhances the progressive strengthening of cortico-cortical connectivity across individual sleep spindles, whereas TMR, at least in the present experimental conditions, attenuates this learning-related evolution. Importantly, the absence of a direct behavioral association indicates that this temporal evolution should not necessarily be interpreted as a direct marker of overnight performance gains. Rather, it may represent one component of a broader network-level reorganization during NREM sleep, potentially interacting with microstructural plasticity and cortico-subcortical communication during memory consolidation. Notably, TMR altered both spindle occurrence and the temporal dynamics of spindle-related connectivity despite the absence of a detectable behavioral benefit. This suggests that acoustic stimulation during sleep can modify the organization of sleep-related neural activity without necessarily producing measurable changes in next-day performance.

### 4.6. Limitations

A few limitations should be considered when interpreting the present findings. First, the comparison between the habituation and experimental nights may be influenced by a first-night effect. However, the relatively stable sleep architecture observed across nights in the RS group suggests that such effect was likely limited, although it cannot be completely excluded.

Second, the study did not include a third group performing a control task. Consequently, we cannot fully disentangle effects specifically related to motor sequence learning from changes associated with task performance or other non-specific aspects of the experimental procedure. Nevertheless, the comparison between the RS and TMR groups allowed us to isolate the additional effect of TMR on sleep oscillations and connectivity following the same learning experience.

Third, our connectivity analysis was restricted to the cortical surface, as constrained by the spatial resolution and source-reconstruction properties of hd-EEG, as well as ongoing concerns regarding the reliability of deep-source estimation [88]. Consequently, our analysis did not capture connectivity involving deeper subcortical structures, which would have been particularly relevant given the proposed contribution of thalamic, striatal, and hippocampal regions to spindle-related memory processing. The absence of deep-source connectivity therefore limits our ability to characterize the full cortico-subcortical network underlying spindle-related memory processing and to directly compare our findings with studies that have estimated subcortical connectivity, such as Boutin et al. [6]. Future studies combining cortical and validated subcortical source estimates could provide a more comprehensive characterization of these network dynamics.

Finally, methodological differences across studies—including task design, sleep-stage restrictions, spindle detection and classification methods, stimulation protocols, and sample characteristics—may contribute to variability in reported effects. These differences should therefore be considered when comparing the present findings with other TMR and spindle- connectivity studies.

### 4.7. Future directions

The present findings also raise several directions for future research. First, the use of shorter and more temporally constrained auditory cues may help minimize disruption of endogenous spindle and SO dynamics while still providing effective memory reactivation. However, optimizing cue timing may be more complex than simply targeting a fixed interval after the preceding spindle. Recent evidence suggests that spindle occurrence follows highly individualized temporal patterns, including a refractory period followed by a period of increased spindle probability, with these temporal signatures being relatively stable within individuals across nights [113]. This raises the possibility that future TMR approaches could benefit from individualized, real-time stimulation protocols that account for the ongoing dynamics of each participant’s spindle activity rather than relying on fixed stimulation intervals.

A second important direction would be to examine intra-spindle connectivity specifically for SO-coupled versus uncoupled spindles, especially when synchronizing with the SO upstates. This could help determine whether the connectivity dynamics observed in the present study represent a general feature of spindle activity or whether coupled spindles express a different connectivity pattern, thought to be most favorable for memory processing and plasticity.

Finally, future studies combining high-density EEG with more reliable estimates of subcortical activity, alongside longitudinal diffusion imaging, could provide a more complete account of how learning-related structural changes interact with spindle-related network communication. Such approaches could help determine whether the temporal evolution of spindle connectivity represents a mechanism through which newly acquired motor memories are progressively redistributed across hippocampal, striatal, thalamic, cerebellar, and cortical networks.

## 5. Conclusions

In conclusion, the present findings provide evidence that sleep-dependent motor memory consolidation is accompanied by coordinated changes across multiple levels of brain organization, from microstructural properties to sleep oscillations and within-spindle network dynamics. Learning-related microstructural changes were associated with subsequent spindle, SO, and SO–spindle activity, while motor learning also enhanced the progressive strengthening of cortico-cortical connectivity across individual spindles. These findings suggest that the neural consequences of learning may shape the sleep environment in which newly acquired memories are subsequently processed and reorganized.

TMR, however, did not enhance these processes under the present stimulation conditions. Instead, auditory stimulation was associated with reduced spindle and SO activity, diminished SO–spindle co-occurrence, and attenuated intra-spindle connectivity evolution, despite no detectable behavioral impairment. This dissociation between changes in sleep neurophysiology and overt memory performance highlights the complexity of the relationship between sleep oscillations and memory consolidation, and suggests that modifying sleep activity does not necessarily translate into measurable behavioral effects.

More broadly, the present results support a view of sleep-dependent motor memory consolidation as a dynamic interaction between learning-induced structural plasticity and temporally organized neural activity during sleep. Understanding how these processes interact, and how they can be selectively manipulated without disrupting endogenous sleep dynamics, will be important for elucidating the mechanisms underlying memory consolidation and for developing more precise TMR approaches.

## 6. List of abbreviations

AASM: American Academy of Sleep Medicine
ANOVA: Analysis of variance
BMI: Body mass index
dMRI: Diffusion magnetic resonance imaging
DTI: Diffusion tensor imaging
DWI: Diffusion-weighted imaging
EEG: Electroencephalography
EMG: Electromyography
EOG: Electrooculography
FA: Fractional anisotropy
FWF: Free water fraction
GM: Grey matter
GPI: Global Performance Index
hd-EEG: High-density electroencephalography
MD: Mean diffusivity
MEM: Maximum Entropy on the Mean
MNI: Montreal Neurological Institute
MRI: Magnetic resonance imaging
N2/N3: Non-rapid eye movement sleep stages 2 and 3
NBS: Network-Based Statistic
NDI: Neurite density index
NODDI: Neurite orientation dispersion and density imaging
NREM: Non-rapid eye movement
ODI: Orientation dispersion index
PVT: Psychomotor Vigilance Test
ROI: Region of interest
RS: Regular sleep
SD: Standard deviation
SO: Slow oscillation
SRTT: Serial Reaction Time Task
SWA: Slow-wave activity
TMR: Targeted memory reactivation
WM: White matter
wPLI: Weighted Phase Lag Index

## 7. Declarations

### 7.1. Ethics approval and consent to participate

The study was conducted in accordance with the Declaration of Helsinki and was approved by the local Ethics Committee of the Liège University Hospital (approval no. 2020/138). Written informed consent was obtained from all participants prior to participation.

### 7.2. Consent for publication

Not applicable.

### 7.3. Availability of data and materials

The data underlying this article are available upon reasonable request to the corresponding author. The material generated and/or analysed during the current study are available in the *HD-EEG-Spindle-Connectivity* repository (https://github.com/jmarclina/HD-EEG-Spindle-Connectivity/tree/main).

## 7.6. Authors’ contributions

Conceptualization: W.S. and P.P.; Methodology: W.S. J-M.L., and P.P.; Data collection: W.S.; Data analysis: W.S., M-C.F., A.L., M.G., H.Z., J-M.L., and P.P.; Visualization: W.S., J-M.L., and P.P.; Writing—original draft preparation: W.S.; Writing—review and editing: P.P., M-C.F., and J-M.L.; Supervision: P.P., J-M.L., and H.Z.; Funding acquisition: P.P. All authors read and approved the final manuscript.

## Acknowledgements

The authors thank Dimitri Voisin and Louis-Jacques Etaix for their help in data acquisition.

