## Additional File for "Modulating Sleep Oscillations Through Targeted Memory Reactivation: Linking Functional Dynamics to Brain Microstructure"

#### 1. Connectivity Analyses

The methodological framework comprises four main components: 1) the definition of the trials and the detection of grapho-elements, namely sleep spindles and slow waves; 2) the selection and characterization of the spindles events; 3) the source localization of coincident spindles; and 4) the assessment of intra-spindle connectivity. The overall methodology follows the approach presented in [1]. Here, we focus on the main methodological differences that enabled this approach to be adapted to the present study. More specifically, the methodology is structured around the following steps:

- *Trials and the detection pipeline*

Trials are extracted from artefact-free hd-EEG recordings acquired during the two nights and the two first cycles. Each trial consists of a 4096-sample segment (sampling frequency = 250Hz) recorded from 256 sensors. For each trial, information regarding the night, sleep cycles and sleep stage is retained. When applicable, stimulation intervals and associated markers are also stored. Spindle detection was based on the approach proposed by Mölle et al. [2], applied across all EEG sensors. Spindles outside the stimulations are kept in the study. Slow-wave detection followed the approach proposed by Carrier et al. [3] according to standard parameters outlined in [4].

- *The definition of the spindle events*

To increase the specificity of the automatic detection, we retained spindles that were detected concurrently on at least 10% of the sensors, while excluding events detected on more than 40% of the sensors. Coincidence is defined over 500 msec. This selection criterion increases our confidence in source localization in the absence of a subject-specific forward model. The Hilbert transform (10-16Hz) of the segments corresponding to each selected spindle yields complex-valued signals over the sensors suitable for source localization [5].

- *The source localization pipeline*

The Maximum Entropy on the Mean (MEM) approach is a Bayesian inference method introduced and evaluated in [6]. We essentially reproduced the approach described in [1], adapting it to hd-EEG recordings through the use of an appropriate forward model, and to the description of source activity using the Fourier modes associated with the graphs defined on the cortical mesh. The Fourier modes are computed for each anatomical region according to the Destrieux atlas (N = 148 cortical regions). This graphical modeling framework provides a stronger justification for: 1) the use of a common anatomical template of cortical sources across subjects; and 2) a Bernoulli–Gaussian mixture defined the Bayesian prio**r**, with a specific and data-driven prior variance assigned to each 'active' Fourier mode [7].

- *The intra-spindle connectivity pipeline.*

The localization of each spindle provides the time courses of the real and imaginary components of the estimated sources. These complex-valued source signals are first averaged within each of the atlas regions and subsequently divided into two equal-duration segments corresponding to the onset and offset phases of the spindle. For each segment, the resulting N complex-valued regional signals Si(k), i = 1,2,3…,N are used to estimate inter-regional functional connectivity using the weighted Phase Lag Index (wPLI).

$$C_{ij} = abs(E[Im(S_i(k)\ S_j(k)^*)]) / (E[abs(Im(S_i(k)\ S_j(k)^*)]), \quad i,j = 1,2,3,... N.$$

### 2. Figures

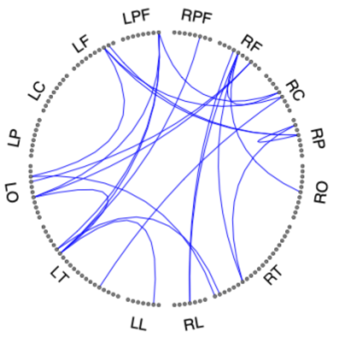


*Additional Figure 1.* RS group - Night 1: synthetic Destrieux connectome associated with wPLI connectivity metrics, generated using the LiveScript 'HD-connectome' Toolbox.

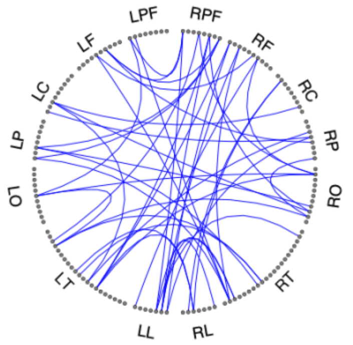


*Additional Figure 2.* TMR group - Night 1: synthetic Destrieux connectome associated with wPLI connectivity metrics, generated using the LiveScript 'HD-connectome' Toolbox.

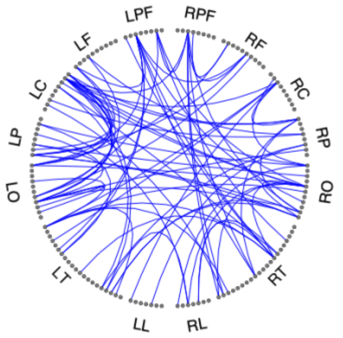


*Additional Figure 3.* RS group - Night 2: synthetic Destrieux connectome associated with wPLI connectivity metrics, generated using the LiveScript 'HD-connectome' Toolbox.

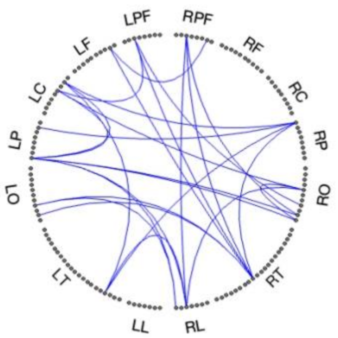


*Additional Figure 4.* RS group - Night 1 vs Night 2: synthetic Destrieux connectome associated with higher proportional connectivity changes in wPLI connectivity metrics in Night 2 than in Night 1, generated using the LiveScript 'HD-connectome' Toolbox.

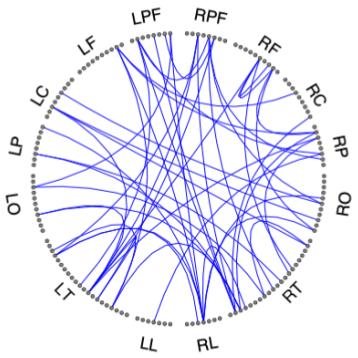


*Additional Figure 5.* TMR group - Night 1 vs Night 2: synthetic Destrieux connectome associated with higher proportional connectivity changes in wPLI connectivity metrics in Night 1 than in Night 2, generated using the LiveScript 'HD-connectome' Toolbox.

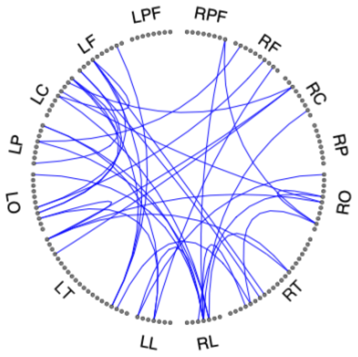


*Additional Figure 6.* Experimental Night 2: RS vs TMR group: synthetic Destrieux connectome associated with higher proportional connectivity changes in wPLI connectivity metrics in the RS than in the TMR group during Night 2, generated using the LiveScript 'HD-connectome' Toolbox

### 3. Tables

#### ADDITIONAL TABLE 1. RS group – Baseline Night 1

| X | Y | Z | Node strength | Label |
|---|---|---|---|---|
| | | | NETWORK 1 | |
| 45 | -2 | 37 | 0,053 | S_precentral-inf-partR |
| 20 | 28 | -15 | 0,041 | S_orbital-H_ShapedR |
| 5 | -37 | 62 | 0,039 | G_and_S_paracentralR |
| -47 | -10 | 28 | 0,039 | G_and_S_subcentralL |
| 38 | -6 | 3 | 0,037 | S_circular_insula_infR |
| 31 | -75 | -10 | 0,03 | S_occipital_antR |
| Best | | | | |
| -6 | -24 | 47 | 0,028 | S_cingul-MarginalisL |
| -8 | 16 | -11 | 0,025 | G_subcallosalL |
| -41 | 15 | -33 | 0,024 | Pole_temporalL |
| -20 | 28 | -15 | 0,021 | S_orbital-H_ShapedL |
| -23 | 29 | -11 | 0,02 | S_orbital_lateralL |
| 17 | -96 | 15 | 0,019 | S_oc_sup_and_transversalR |
| -11 | 28 | -13 | 0,019 | S_orbital_med-olfactL |
| 50 | -2 | -4 | 0,013 | G_temp_sup-Plan_polarR |
| -45 | -2 | 37 | 0,012 | S_precentral-inf-partL |
| 4 | 30 | 30 | 0,01 | G_and_S_cingul-Mid-AntR |
| -3 | -33 | 41 | 0,01 | G_cingul-Post-dorsalL |
| 6 | -16 | 25 | 0,01 | S_pericallosalR |
| 24 | 52 | -3 | 0,01 | G_and_S_transv_frontopolR |
| 43 | 13 | 16 | 0,01 | G_front_inf-OpercularR |
| -9 | -50 | 31 | 0,009 | S_subparietalL |
| 55 | -28 | 11 | 0,009 | G_temp_sup-Plan_tempoR |
| 31 | 36 | 22 | 0,009 | S_front_middleR |
| 46 | -48 | -12 | 0,008 | S_oc-temp_latR |
| -28 | -91 | -8 | 0,008 | G_and_S_occipital_infL |
| -24 | -47 | 49 | 0,007 | S_intrapariet_and_P_transL |
| | | | NETWORK 2 | |
| 20 | 52 | 26 | 0,047 | G_front_supR |
| -38 | 5 | 11 | 0,047 | S_circular_insula_supL |
| -20 | 43 | -18 | 0,047 | S_suborbitalL |
| -20 | -89 | 29 | 0,034 | G_occipital_supL |
| 8 | -56 | 43 | 0,03 | G_precuneusR |
| -41 | 22 | -10 | 0,029 | S_front_infL |
| 41 | -16 | 49 | 0,025 | S_centralR |

| | | | | |
|---|---|---|---|---|
| -35 | -27 | -11 | 0,021 | S_collat_transv_antL |
| 3 | -33 | 41 | 0,02 | G_cingul-Post-dorsalR |
| 38 | 5 | 11 | 0,02 | S_circular_insula_supR |
| 41 | 22 | -10 | 0,02 | S_front_infR |
| -21 | -90 | 9 | 0,02 | S_oc_middle_and_LunatusL |
| 6 | -24 | 47 | 0,019 | S_cingul-MarginalisR |
| 50 | -45 | 44 | 0,015 | G_pariet_inf-SupramarR |
| -4 | 30 | 30 | 0,014 | G_and_S_cingul-Mid-AntL |
| -31 | 36 | 22 | 0,012 | S_front_middleL |
| 18 | -100 | 2 | 0,011 | Pole_occipitalR |
| -27 | -38 | -12 | 0,011 | S_collat_transv_postL |
| 49 | -28 | 11 | 0,011 | S_temporal_transverseR |
| -47 | -20 | 8 | 0,01 | G_temp_sup-G_T_transvL |
| 52 | -18 | 0 | 0,01 | S_temporal_supR |
| 34 | 40 | 28 | 0,01 | G_front_middleR |
| 23 | 29 | -11 | 0,008 | S_orbital_lateralR |

*Additional Table 1.* RS group - Night 1: List of nodes of the significant networks in which wPLI connectivity evolved from the beginning to the end of the spindles in sleep stages N2-N3 during the second sleep cycle. MNI coordinates (X, Y, Z) are reported in millimetres (mm).

**ADDITIONAL TABLE 2. TMR group – Baseline Night 1**

| X | Y | Z | Node strength | Label |
|---|---|---|---|---|
| -50 | -2 | -4 | 0,05 | G_temp_sup-Plan_polarL |
| -4 | 30 | 30 | 0,044 | G_and_S_cingul-Mid-AntL |
| 2 | -83 | 17 | 0,041 | G_cuneusR |
| 52 | -18 | 0 | 0,04 | S_temporal_supR |
| 3 | -38 | 24 | 0,034 | G_cingul-Post-ventralR |
| 23 | 29 | -11 | 0,034 | S_orbital_lateralR |
| -38 | -26 | 52 | 0,033 | G_postcentralL |
| -33 | 21 | -8 | 0,029 | Lat_Fis-ant-VerticalL |
| -18 | -100 | 2 | 0,029 | Pole_occipitalL |
| 26 | 59 | 5 | 0,028 | G_and_S_frontomarginR |
| -14 | -65 | -5 | 0,028 | S_oc-temp_med_and_LingualL |
| 20 | 43 | -18 | 0,027 | S_suborbitalR |
| -5 | -19 | 36 | 0,026 | G_and_S_cingul-Mid-PostL |
| -49 | -15 | -34 | 0,026 | G_temporal_infL |
| -23 | -56 | 51 | 0,025 | G_parietal_supL |
| 8 | -56 | 43 | 0,023 | G_precuneusR |
| 7 | 34 | -15 | 0,023 | G_rectusR |
| 21 | -90 | 9 | 0,022 | S_oc_middle_and_LunatusR |
| -8 | 34 | 18 | 0,021 | G_and_S_cingul-AntL |
| -26 | 59 | 5 | 0,021 | G_and_S_frontomarginL |
| 41 | 28 | 17 | 0,021 | G_front_inf-TriangulR |
| -43 | -63 | 33 | 0,02 | G_pariet_inf-AngularL |
| 33 | 21 | -8 | 0,02 | Lat_Fis-ant-VerticalR |
| 47 | -10 | 28 | 0,019 | G_and_S_subcentralR |
| 17 | -96 | 15 | 0,019 | S_oc_sup_and_transversalR |
| 54 | -20 | -22 | 0,019 | S_temporal_infR |
| 43 | 13 | 16 | 0,018 | G_front_inf-OpercularR |
| -20 | 52 | 26 | 0,018 | G_front_supL |
| 34 | 10 | 0 | 0,017 | S_circular_insula_antR |
| 23 | -56 | 51 | 0,013 | G_parietal_supR |
| 8 | 34 | 18 | 0,011 | G_and_S_cingul-AntR |
| 26 | -46 | 40 | 0,011 | S_interm_prim-JensenR |

| | | | | |
|---|---|---|---|---|
| -6 | -16 | 25 | 0,011 | S_pericallosalL |
| 9 | -50 | 31 | 0,011 | S_subparietalR |
| -54 | -20 | -22 | 0,011 | S_temporal_infL |
| -24 | 52 | -3 | 0,01 | G_and_S_transv_frontopolL |
| -41 | 15 | -33 | 0,01 | Pole_temporalL |
| 8 | 16 | -11 | 0,009 | G_subcallosalR |
| -5 | -37 | 62 | 0,009 | G_and_S_paracentralL |
| 34 | 15 | 6 | 0,009 | G_insular_shortR |
| 45 | -2 | 37 | 0,009 | S_precentral-inf-partR |
| 38 | -6 | 3 | 0,009 | S_circular_insula_infR |
| 39 | 14 | -8 | 0,008 | Lat_Fis-ant-HorizontR |
| -50 | -35 | 11 | 0,008 | Lat_Fis-postL |
| 27 | -38 | -12 | 0,008 | S_collat_transv_postR |
| -47 | -20 | 8 | 0,01 | G_temp_sup-G_T_transvL |
| 52 | -18 | 0 | 0,01 | S_temporal_supR |
| 34 | 40 | 28 | 0,01 | G_front_middleR |
| 23 | 29 | -11 | 0,008 | S_orbital_lateralR |

*Additional Table 2.* TMR group - Night 1: List of nodes of the significant networks in which wPLI connectivity evolved from the beginning to the end of the spindles in sleep stages N2-N3 during the second sleep cycle. MNI coordinates (X, Y, Z) are reported in millimetres (mm).

**ADDITIONAL TABLE 3. RS group – Experimental Night 2**

| X | Y | Z | Node strength | Label |
|---|---|---|---|---|
| -45 | -2 | 37 | 0,097 | S_precentral-inf-partL |
| 20 | -89 | 29 | 0,094 | G_occipital_supR |
| -32 | -82 | 8 | 0,083 | G_occipital_middleL |
| -21 | 28 | -13 | 0,08 | G_orbitalL |
| 21 | 28 | -13 | 0,075 | G_orbitalR |
| 55 | -28 | 11 | 0,064 | G_temp_sup-Plan_tempoR |
| -49 | -28 | 11 | 0,063 | S_temporal_transverseL |
| -43 | -63 | 33 | 0,061 | G_pariet_inf-AngularL |
| -31 | -75 | -10 | 0,055 | S_occipital_antL |
| 31 | -75 | -10 | 0,051 | S_occipital_antR |
| 43 | -63 | 33 | 0,05 | G_pariet_inf-AngularR |
| 38 | -26 | 52 | 0,049 | G_postcentralR |
| -26 | -18 | 63 | 0,045 | S_precentral-sup-partL |
| 17 | -96 | 15 | 0,044 | S_oc_sup_and_transversalR |
| -11 | 28 | -13 | 0,042 | S_orbital_med-olfactL |
| 28 | -91 | -8 | 0,035 | G_and_S_occipital_infR |
| 3 | -38 | 24 | 0,034 | G_cingul-Post-ventralR |
| -20 | -89 | 29 | 0,033 | G_occipital_supL |
| 34 | 10 | 0 | 0,031 | S_circular_insula_antR |
| -43 | 13 | 16 | 0,03 | G_front_inf-OpercularL |
| 39 | 14 | -8 | 0,027 | Lat_Fis-ant-HorizontR |
| 5 | -19 | 36 | 0,023 | G_and_S_cingul-Mid-PostR |
| -22 | -36 | -16 | 0,023 | G_oc-temp_med-ParahipL |
| 34 | 33 | -7 | 0,022 | G_front_inf-OrbitalR |
| -34 | 40 | 28 | 0,021 | G_front_middleL |
| -23 | -56 | 51 | 0,02 | G_parietal_supL |
| 52 | -18 | 0 | 0,02 | S_temporal_supR |
| 8 | -56 | 43 | 0,019 | G_precuneusR |
| -47 | -20 | 8 | 0,019 | G_temp_sup-G_T_transvL |
| 24 | -47 | 49 | 0,019 | S_intrapariet_and_P_transR |
| 49 | -28 | 11 | 0,019 | S_temporal_transverseR |
| -39 | -41 | -20 | 0,018 | G_oc-temp_lat-fusiforL |

| | | | | |
|---|---|---|---|---|
| 50 | -2 | -4 | 0,017 | G_temp_sup-Plan_polarR |
| -6 | -24 | 47 | 0,017 | S_cingul-MarginalisL |
| -38 | 5 | 11 | 0,017 | S_circular_insula_supL |
| -16 | -72 | 18 | 0,017 | S_parieto_occipitalL |
| -35 | -27 | -11 | 0,012 | S_collat_transv_antL |
| -2 | -83 | 17 | 0,011 | G_cuneusL |
| 14 | -62 | -3 | 0,011 | G_oc-temp_med-LingualR |
| 7 | 34 | -15 | 0,011 | G_rectusR |
| 18 | -100 | 2 | 0,011 | Pole_occipitalR |
| -14 | -65 | -5 | 0,011 | S_oc-temp_med_and_LingualL |
| 16 | -72 | 18 | 0,011 | S_parieto_occipitalR |
| 54 | -20 | -22 | 0,011 | S_temporal_infR |
| -5 | -19 | 36 | 0,01 | G_and_S_cingul-Mid-PostL |
| -38 | -26 | 52 | 0,01 | G_postcentralL |
| 33 | 21 | -8 | 0,01 | Lat_Fis-ant-VerticalR |
| -27 | -38 | -12 | 0,01 | S_collat_transv_postL |
| 27 | -38 | -12 | 0,01 | S_collat_transv_postR |
| -39 | -27 | 46 | 0,01 | S_postcentralL |
| 47 | -10 | 28 | 0,01 | G_and_S_subcentralR |
| -47 | -10 | 28 | 0,01 | G_and_S_subcentralL |
| -50 | -45 | 44 | 0,01 | G_pariet_inf-SupramarL |
| -26 | 59 | 5 | 0,01 | G_and_S_frontomarginL |
| -55 | -28 | -12 | 0,01 | G_temporal_middleL |
| -14 | -62 | -3 | 0,009 | G_oc-temp_med-LingualL |
| -26 | -46 | 40 | 0,009 | S_interm_prim-JensenL |
| -34 | 15 | 6 | 0,009 | G_insular_shortL |
| 22 | -36 | -16 | 0,009 | G_oc-temp_med-ParahipR |
| 6 | -16 | 25 | 0,008 | S_pericallosalR |
| -6 | -16 | 25 | 0,008 | S_pericallosalL |

*Additional Table 3.* RS group - Night 2: List of nodes of the significant networks in which wPLI connectivity evolved from the beginning to the end of the spindles in sleep stages N2-N3 during the second sleep cycle. MNI coordinates (X, Y, Z) are reported in millimetres (mm).

**ADDITIONAL TABLE 4. RS group – Baseline Night 1 vs Experimental Night 2**

| X | Y | Z | Node strength | Label |
|---|---|---|---|---|
| 34 | 10 | 0 | 17,00 | S_circular_insula_antR |
| -43 | -63 | 33 | 16,00 | G_pariet_inf-AngularL |
| 3 | -38 | 24 | 13,00 | G_cingul-Post-ventralR |
| 43 | -63 | 33 | 11,00 | G_pariet_inf-AngularR |
| 20 | -89 | 29 | 9,70 | G_occipital_supR |
| -22 | -36 | -16 | 9,50 | G_oc-temp_med-ParahipL |
| -45 | -2 | 37 | 9,10 | S_precentral-inf-partL |
| 21 | 28 | -13 | 8,50 | G_orbitalR |
| -21 | 28 | -13 | 8,10 | G_orbitalL |
| -6 | -24 | 47 | 7,60 | S_cingul-MarginalisL |
| 31 | -75 | -10 | 6,10 | S_occipital_antR |
| -32 | -82 | 8 | 5,70 | G_occipital_middleL |
| 6 | -16 | 25 | 5,60 | S_pericallosalR |
| -31 | 36 | 22 | 5,30 | S_front_middleL |
| 17 | -96 | 15 | 5,20 | S_oc_sup_and_transversalR |
| -16 | -72 | 18 | 3,20 | S_parieto_occipitalL |
| 27 | -38 | -12 | 2,70 | S_collat_transv_postR |
| -2 | -83 | 17 | 2,30 | G_cuneusL |
| 11 | 28 | -13 | 2,30 | S_orbital_med-olfactR |

*Additional Table 4.* RS group - Night 1 vs Night 2 : List of nodes of the significant networks in which proportional connectivity changes from spindles beginning to end (sleep stages N2-N3 during the second sleep cycle) were higher in Experimental Night 2 than in Baseline Night 1. MNI coordinates (X, Y, Z) are reported in millimetres (mm).

**ADDITIONAL TABLE 5. TMR group – Baseline Night 1 vs Experimental Night 2**

| X | Y | Z | Node strength | Label |
|---|---|---|---|---|
| 3 | -33 | 41 | -21,00 | G_cingul-Post-dorsalR |
| -49 | -15 | -34 | -20,00 | G_temporal_infL |
| -24 | 52 | -3 | -16,00 | G_and_S_transv_frontopolL |
| -31 | 36 | 22 | -14,00 | S_front_middleL |
| 39 | 14 | -8 | -13,00 | Lat_Fis-ant-HorizontR |
| 20 | 28 | -15 | -13,00 | S_orbital-H_ShapedR |
| 52 | -18 | 0 | -13,00 | S_temporal_supR |
| 54 | -20 | -22 | -10,00 | S_temporal_infR |
| 50 | -45 | 44 | -9,90 | G_pariet_inf-SupramarR |
| -38 | -26 | 52 | -9,90 | G_postcentralL |
| 21 | 28 | -13 | -9,60 | G_orbitalR |
| -22 | -36 | -16 | -9,20 | G_oc-temp_med-ParahipL |
| 31 | 36 | 22 | -9,10 | S_front_middleR |
| -14 | -65 | -5 | -8,80 | S_oc-temp_med_and_LingualL |
| 24 | -47 | 49 | -7,40 | S_intrapariet_and_P_transR |
| 18 | -100 | 2 | -6,90 | Pole_occipitalR |
| -50 | -2 | -4 | -6,80 | G_temp_sup-Plan_polarL |
| -20 | 43 | -18 | -6,80 | S_suborbitalL |
| 23 | 29 | -11 | -6,40 | S_orbital_lateralR |
| -21 | -90 | 9 | -6,30 | S_oc_middle_and_LunatusL |
| 34 | 15 | 6 | -6,20 | G_insular_shortR |
| 23 | -56 | 51 | -6,00 | G_parietal_supR |
| -23 | -56 | 51 | -5,80 | G_parietal_supL |
| 38 | -6 | 3 | -5,70 | S_circular_insula_infR |
| 31 | -75 | -10 | -5,50 | S_occipital_antR |
| -32 | -82 | 8 | -5,30 | G_occipital_middleL |
| 50 | -2 | -4 | -3,70 | G_temp_sup-Plan_polarR |
| 41 | -6 | 48 | -3,60 | G_precentralR |
| 41 | 15 | -33 | -3,50 | Pole_temporalR |
| 21 | -90 | 9 | -3,50 | S_oc_middle_and_LunatusR |
| -6 | -24 | 47 | -3,40 | S_cingul-MarginalisL |

| | | | | |
|---|---|---|---|---|
| -50 | -35 | 11 | -3,20 | Lat_Fis-postL |
| 55 | -28 | -12 | -3,10 | G_temporal_middleR |
| 41 | 22 | -10 | -3,10 | S_front_infR |
| -54 | -20 | -22 | -3,10 | S_temporal_infL |
| 3 | -38 | 24 | -3,00 | G_cingul-Post-ventralR |
| -20 | 28 | -15 | -3,00 | S_orbital-H_ShapedL |
| -9 | -50 | 31 | -3,00 | S_subparietalL |
| -3 | -33 | 41 | -2,90 | G_cingul-Post-dorsalL |
| 14 | -65 | -5 | -2,80 | S_oc-temp_med_and_LingualR |
| -11 | 28 | -13 | -2,80 | S_orbital_med-olfactL |
| 16 | -72 | 18 | -2,60 | S_parieto_occipitalR |
| 26 | -18 | 63 | -2,50 | S_precentral-sup-partR |
| 20 | -89 | 29 | -2,40 | G_occipital_supR |
| 55 | -28 | 11 | -2,30 | G_temp_sup-Plan_tempoR |

*Additional Table 5.* TMR group - Night 1 vs Night 2 : List of nodes of the significant networks in which proportional connectivity changes from spindles beginning to end (sleep stages N2-N3 during the second sleep cycle) were higher in Baseline Night 1 than in Experimental Night 2. MNI coordinates (X, Y, Z) are reported in millimetres (mm).

**ADDITIONAL TABLE 6. Experimental Night 2: RS vs TMR group**

| X | Y | Z | Node strength | Label |
|---|---|---|---|---|
| 3 | -33 | 41 | -20,00 | G_cingul-Post-dorsalR |
| -34 | 40 | 28 | -14,00 | G_front_middleL |
| -49 | -28 | 11 | -14,00 | S_temporal_transverseL |
| 5 | -19 | 36 | -11,00 | G_and_S_cingul-Mid-PostR |
| -22 | -36 | -16 | -11,00 | G_oc-temp_med-ParahipL |
| 38 | -26 | 52 | -11,00 | G_postcentralR |
| -26 | -18 | 63 | -10,00 | S_precentral-sup-partL |
| -3 | -33 | 41 | -9,50 | G_cingul-Post-dorsalL |
| 17 | -96 | 15 | -8,40 | S_oc_sup_and_transversalR |
| -26 | -46 | 40 | -8,10 | S_interm_prim-JensenL |
| -6 | -16 | 25 | -7,50 | S_pericallosalL |
| -20 | -89 | 29 | -7,30 | G_occipital_supL |
| 18 | -100 | 2 | -7,20 | Pole_occipitalR |
| -41 | -16 | 49 | -7,10 | S_centralL |
| -9 | -50 | 31 | -7,10 | S_subparietalL |
| 55 | -28 | 11 | -6,80 | G_temp_sup-Plan_tempoR |
| -43 | 13 | 16 | -6,50 | G_front_inf-OpercularL |
| -14 | -62 | -3 | -6,10 | G_oc-temp_med-LingualL |
| 46 | -48 | -12 | -6,00 | S_oc-temp_latR |
| -32 | -82 | 8 | -5,80 | G_occipital_middleL |
| 20 | 43 | -18 | -5,80 | S_suborbitalR |
| 20 | -89 | 29 | -5,50 | G_occipital_supR |
| -31 | 36 | 22 | -3,90 | S_front_middleL |
| -14 | -65 | -5 | -3,90 | S_oc-temp_med_and_LingualL |
| 34 | 33 | -7 | -3,80 | G_front_inf-OrbitalR |
| -43 | -63 | 33 | -3,80 | G_pariet_inf-AngularL |
| 38 | 5 | 11 | -3,70 | S_circular_insula_supR |
| 33 | 21 | -8 | -3,60 | Lat_Fis-ant-VerticalR |
| -6 | -24 | 47 | -3,60 | S_cingul-MarginalisL |
| 45 | -2 | 37 | -3,50 | S_precentral-inf-partR |
| 28 | -91 | -8 | -3,30 | G_and_S_occipital_infR |

| | | | | |
|---|---|---|---|---|
| -45 | -2 | 37 | -3,30 | S_precentral-inf-partL |
| -39 | -41 | -20 | -3,20 | G_oc-temp_lat-fusiforL |
| 31 | 36 | 22 | -3,10 | S_front_middleR |
| 14 | -65 | -5 | -3,00 | S_oc-temp_med_and_LingualR |
| 3 | -38 | 24 | -2,90 | G_cingul-Post-ventralR |
| 38 | -6 | 3 | -2,90 | S_circular_insula_infR |
| -31 | -75 | -10 | -2,90 | S_occipital_antL |

*Additional Table 6.* Experimental Night 2: RS vs TMR group: List of nodes of the significant networks in which proportional connectivity changes from spindles beginning to end (sleep stages N2-N3 during the second sleep cycle) were higher in the RS than the TMR group during Experimental Night 2. MNI coordinates (X, Y, Z) are reported in millimetres (mm).
